# Quantification of extracellular vesicles with unaltered surface membranes using an internalized oligonucleotide tracer and droplet digital PCR applied to modeling of *in vivo* kinetics

**DOI:** 10.1101/2020.09.22.306969

**Authors:** Thomas De Luca, Robert E. Stratford, Madison E. Edwards, Christina R. Ferreira, Eric A. Benson

## Abstract

Extracellular vesicles (EVs) continue to attract interest for their potential role in targeted therapeutics and as biomarkers of disease and drug response. In order to achieve clinical utility, it is important to determine the pharmacokinetic parameters of candidate EVs during preclinical development using *in vivo* animal models. To date, no methods exist for studying EV kinetics without modification to surface ligands that may affect normal behavior. Here we introduce an accessible method for labeling and quantifying EVs administered to conscious animals, without disrupting endogenous ligands. Our method relies upon established laboratory techniques and can be tailored to a variety of biological questions. Digital PCR is leveraged to detect a non-homologous oligonucleotide tracer introduced into the vesicles, allowing for quantification over a wide dynamic range. Using an application of this method, we found differences in the *in vivo* kinetics of EVs from three different cell types using non-linear mixed effects modeling. We propose that this method will provide a complementary approach for the of study EV ligand-receptor interactions in the context of EV uptake and targeted therapeutics.

## Introduction

Extracellular vesicles (EV) can be used to improve medical treatments if properly understood^1,2^. Chief among the EV subtypes that have captured the interest of clinical researchers are exosomes, which are small (<200 nm) EVs that begin as the intraluminal vesicles of the late stage endosome, where they are loaded with active biological molecules such as microRNAs (miRNA), mRNA, and proteins^3^. Once secreted, they transport these contents to other nearby cells or to distant tissues via the blood circulation. Targeted distribution of these vesicles is governed by surface markers, the composition of which is dependent on the originating cell^4-6^. Since EVs are continually secreted by virtually every eukaryotic cell, it is broadly accepted that the composition of any individual vesicle reflects the status of its originating cell at a particular moment in time. This dynamic heterogeneity in blood-circulating EVs makes the study of EV kinetics difficult^7,8^.

In order to quantitatively decipher the complexity of circulating EVs, there is a need for an easily applicable, reproducible method for determining the kinetic parameters of EVs from known origins^2^. Due to the inherent difficulty of studying EV transport and distribution in humans, preclinical *in vivo* animal models are used. Existing studies of circulating EV kinetics are limited and have involved the development of membrane-associated labels and companion detection methods. The use of luciferase or radiolabels anchored to exogenously expressed transmembrane proteins^4,9^ may provide exceptional kinetic information for the evaluation of engineered targeted therapeutics, but it is not ideal for the study of unmodified EVs. To arrive at a better understanding of how endogenous EV composition affects kinetics, we should measure the kinetics of EVs with unmodified surface membranes.

To address this gap in methodology, we sought to develop an accessible and scalable approach that: 1) allows labeling of EVs without membrane surface modification, 2) provides reproducible and quantitative measurements of kinetic parameters, 3) fits within established workflows for the computational modeling of kinetics, and 4) is readily implemented with conventional reagents in a cost-effective manner. Here we describe a method to label the contents of EVs from cultured cell lines and to measure the kinetics of labeled EVs intravenously administered to animals. We applied this method to test a hypothesis that EVs from different non-cancer cell lines exhibit different kinetics *in vivo*. Labeled EVs were isolated from the enriched media of three different species-matched cell lines using a commercial reagent and introduced into the central circulation of conscious animals. Blood from each animal was collected over time, and the plasma fractions were assayed for tracer concentrations. Using principles of pharmacokinetics, we developed kinetic models of EVs from liver, lung, and kidney-derived cell lines, and report significant differences in the kinetic parameters between them. We discuss how a three-compartment non-linear mixed effects model best describes the data and provides evidence that dispositional properties of circulating EVs are sensitive to imparted biological characteristics unique to their source.

## Materials and Methods

### Animals and Housing Conditions

All procedures were conducted in accordance with applicable federal regulations^10,11^ and following the review and approval by the Indiana University-Purdue University Indianapolis (IUPUI) Institutional Animal Care and Use Committee (project 10715, approved 18 June 2014, and project 11299, approved 12 September 2017). Adult male Hsd:Sprague Dawley rats (*n* = 30; male; >350 g; Envigo, Indianapolis, IN, USA) were housed individually or in pairs under standard environmental conditions (22 °C ± 2 °C; 30-70 % relative humidity; 12:12 h light:dark cycle; lights on at 7:00 AM) for ≥ 14 d prior to surgical manipulations. Following surgical implantation of catheters, all animals were individually housed. animals were housed in individually ventilated microisolator shoebox caging with direct contact bedding (Sani-Chip, Envigo, Indianapolis, IN, USA). Food (6.2 % fat, 18.6 % protein, 3.5 % fiber, cat. No. 2018SX, Teklad, Madison, WI, USA) and water were provided ad libitum. The colony was screened quarterly using indirect sentinels for the following pathogens: Kilham rat virus (KRV), rat parvoviruses (RPV, H-1, RMV), rat coronavirus (SDAV), rat theliovirus (RTV), Clostridium piliforme, Mycoplasma pulmonis, and ecto- and endo-parasites. The colony was free of all pathogens during this study.

### Surgery and catheter maintenance

After a two week acclimation period, animals were subjected to surgery for placement of jugular vein catheters^12^. The animals were anesthetized using isoflurane inhalation (5% induction, 2–3% maintenance) and body weight was measured on a digital scale. Before the incision was made, 5-10 mg/kg ketoprofen (Zoetis Inc., Kalamazoo, MI, USA) was subcutaneously administered, hair was shaved from the incision site and the site was scrubbed 3 times with alternating use of povidone-iodine and 70% isopropyl alcohol. A ventral cervical skin incision (1.5 cm) was made right of the midline of the neck slightly anterior to the level of the clavicle. The jugular vein was located by blunt dissection, then isolated and tied off using 4-0 silk suture (cat. no. MV683, Med-Vet International, Mettawa, IL, USA) to occlude blood flow from the head region. During this process, sterile saline was used to keep the area moist. Next, two microdrops of 0.5% lidocaine were applied to the vein to prevent constriction. A small incision was made into the vein using microscissors, and a catheter (polyurethane, 3.5 fr, cat. no. RJVC0612A, Access Technologies, Skokie, IL, USA) filled with bacteriostatic saline was inserted into the vein and threaded 2.5 cm toward the heart. The catheter was secured in place with a second 4-0 silk suture caudal to the insertion site, around the vein. The previously placed cranial suture was used to secure the exterior catheter; this suture was placed between two moveable beads that are included by the manufacturer. After tightening the suture, the beads were slid together to anchor the catheter in place. An aluminum plug was inserted into the catheter after confirming patency. To exteriorize the catheter, an incision was made at the nape of the neck and a trocar was tunneled subcutaneously to the site of the catheter. After passing the catheter through the trocar, both incisions were closed with non-absorbable 4-0 nylon suture (cat. no. #MV-662-V, Med-Vet International), and the exteriorized catheter was secured with a 9 mm Autoclip (cat. no. 205016, MikRon Precision, Gardena, CA, USA) placed between subcutaneous and exterior PE100 dumbbells. A final non-absorbable nylon suture anchored the exterior dumbbell to the skin. After a final check for patency, the catheter was flushed with 0.6 mL bacteriostatic saline and locked with 0.15 mL 4% sodium citrate (4B7867Q, Fenwal Inc., Lake Zurich, IL, USA). Liquid bandage (CVS Health, Woonsocket, RI, USA) was applied to the exteriorized catheter to extend patency^12^. After 2 d of surgical recovery, patency was verified by placing animals into a rodent restrainer (cat. no. 51335, Stoelting Co., Wood Dale, IL, USA), drawing 0.1 mL blood into a discard syringe (cat. no. 309626, BD, Becton Dickinson, Franklin Lakes, NJ, USA), and pulsatile flushing with 0.4 mL bacteriostatic saline. Catheters were flushed by this method every 48 h or less, and locked with 0.15 mL 4% sodium citrate. To minimize backflow of blood into the catheter lumen, a positive pressure flushing technique was used whenever withdrawing a syringe from the catheter.

### Cell culture

Clone 9 hepatocyte (cat. no. CRL-1439), RFL-6 lung fibroblast (cat. no. CCL-192) and RMC mesangial (cat. no. CRL-2573) cells were obtained from American Type Culture Collections (ATCC, Manassas, VA, USA). Upon receipt, cells were passaged 3 times to create cryopreserved stocks. All cell lines were grown per ATCC recommended culture conditions. Base media for each cell line are as follows: Clone 9 cells were cultured in Corning cellgro F-12K medium (cat. no. MT10025CV, Fisher Scientific, Florence, KY, USA) supplemented with 10% fetal bovine serum (FBS, cat. no. S11150, Atlanta Biologicals, Flowery Branch, GA, USA), RFL-6 cells were cultured in Corning cellgro F-12K medium supplemented with 20% FBS, and RMC cells were cultured in ATCC-formulated DMEM (cat. no. MT10017CV, ATCC) supplemented with 15% FBS. Cryopreservation medium was the same as complete growth medium, supplemented with 5% (v/v) DMSO. All cells were grown at 37 °C, 5% CO_2_. Cells were subcultured every 48-72 h, depending on cell density (75-90%) and growth rate of the specific cell line.

All cell lines were authenticated for correct species and verified free of interspecies or mycoplasma contamination by IDEXX (Westbrook, ME, USA). Since these cell lines have not been previously analyzed by IDEXX, they are now considered to be reference cell lines for all future analyses.

### Bacterial transformation with XMIR-NT and plasmid preparation

An amount of 4 ng of XMIRXpress vector containing Non-Targeting miRNA with Xmotif (cat. no. XMIRXP-NT, System Biosciences LLC, Palo Alto, CA, USA) was introduced into 100 μL Stellar competent *E. Coli* cells (Takara Bio USA Inc., Mountain View, CA, USA) by incubation on ice for 30 min, heat shock at 42 °C for 30 s, and replacement on ice for 2 min. After being warmed to room temperature (RT), the final volume was increased to 1 mL using SOC medium (cat. no. ST0215, Takara) pre-warmed to 37 °C. The microcentrifuge tube was wrapped with Parafilm M (Bemis Company Inc., Neenah, WI, USA) and incubated for 40 min at 37 °C on a Thermo Forma orbital shaker (model 420, Thermo Fisher Scientific, Waltham, MA, USA) at 225 RPM. The transformation was diluted 1:10, and 20 μL were spread using 8 to 12 sterile glass beads (cat. no. 710134, MilliporeSigma, Burlington, MA, USA) onto a 10 cm plate (cat. no. FB0875712, Fisher Scientific) containing sterile agar (cat. no. BP1425-500, Fisher Scientific) with Miller’s LB medium (cat. no. 46050CM, Corning Inc., Corning, NY, USA) and 100 μg/mL ampicillin (cat. no. A9626, Teknova, Hollister, CA, USA). After overnight incubation at 37 °C, two colonies were selected for outgrowth in culture tubes (cat. no. 149569C, Fisher Scientific) containing 5 mL autoclaved LB broth (cat. no. J8331L, Quality Biological Inc., Gaithersburg, MD, USA) supplemented with 100 μg/mL ampicillin (cat. no. A5354, Sigma-Aldritch, St. Louis, MO, USA). Liquid cultures were incubated overnight at 37 °C with shaking. Glycerol stocks were prepared by adding 1.6 mL

bacterial culture to screw-top cryovials (cat. no. 430488, Corning) and adding glycerol (cat. no. 327255000, Thermo Fisher Scientific) to a final concentration of 25%. Plasmid DNA was purified from the remaining cultures using a NucleoSpin Plasmid (NoLid) high-pure plasmid mini prep kit (cat. no. 740499.50, Macherey-Nagel, Bethlehem, PA) according to vendor instructions and quantified using the Qubit dsDNA BR Assay (cat. no. Q32850, Thermo Fisher Scientific) with a Qubit Fluorimeter (cat. no. Q32857, Invitrogen, Carlsbad, CA, USA). Plasmid sequence was verified by total plasmid sequencing through the Massachusetts General Hospital Center for Computational & Integrative Biology (MGH CCIB) DNA Core (Cambridge, MA, USA).

For preparation of large amounts of plasmid DNA, a small amount of glycerol stock scraped with a sterile pipet tip was used to prepare a 3 mL starter culture in LB antibiotic selection media, incubated for 12 h at 37 °C with shaking. 1 mL starter culture was transferred to 160 mL LB broth with 100 μg/mL ampicillin in baffled 1 L flasks (cat. no. 25630-1000, Kimble Chase, Vineland, NJ, USA) and incubated for 20 h at 37 °C with vigorous shaking. Plasmid DNA was extracted using the Qiagen HiSpeed Maxi Kit (cat. no. 12662, Qiagen Inc, Valencia, CA, USA) following the vendor’s protocol.

### XMc39 plasmid design and validation

*Caenorhabditis elegans* miR-39-3p (Accession MI0000010, cel-miR-39-3p) stem-loop sequence was retrieved from www.miRBase.org and used to design oligonucleotides for cloning into the XMIRXpress pre-linearized cloning lentivector (cat. no. XMIRXP-Vect, System Biosciences), according to vendor’s instructions (XMIR-c39-top: 5’- GATCCAGCTGATTTCGTCTTGGTAATAAGCTCGTCATTGAGATTATCACCGGGTGTAAATCAGCTTGC-3’, XMIR-c39-bot: 5’- CTAGGCAAGCTGATTTACACCCGGTGATAATCTCAATGACGAGCTTATTACCAAGACGAAATCAGCTG-3’). “Top” and “bottom” oligonucleotides (IDT, Integrated DNA Technologies, Inc., Coralville, IA, USA) were diluted and mixed into a volume of 20 μL low-EDTA TE buffer (cat. no. 11-05-01-13, IDT) with a final concentration of 1 μM each. Annealing was performed in a Veriti thermal cycler (Applied Biosystems, Foster City, CA, USA) as follows. Denaturation was performed at 95 °C for 2 min. Annealing was performed in 4 steps to minimize the formation of secondary structures: 1) Cooling to 63.8 °C over 20 min at a 30% ramp rate, then holding the sample at 63.8 °C for 10 min; 2) Cooling to 46 °C over 20 min at a 30% ramp rate; 3) Cooling to 23 °C at a 100% ramp rate. 1 μL annealed stem-loop oligonucleotide was mixed with 1 μL pre-linearized vector and ligated with T4 DNA ligase (cat. no. M0202, New England BioLabs, Inc., Ipswich, MA, USA), according to vendor’s ligation protocol. 5 μL ligation product was introduced into 50 μL Stbl3 competent *E. Coli* cells (cat. no. C7373-03, Invitrogen) by incubation on ice for 30 min, heat shock at 42 °C for 45 s, and placement on ice for 2 min. 250 μL room-temperature SOC medium was added, and the transformation was incubated for 1 h at 37 °C with shaking on a Thermo Forma orbital shaker at 225 rpm. Three amounts (25 μL, 50 μL, and 100 μL) of the liquid culture were used for antibiotic selection as previously described. 4 colonies were selected and used to prepare glycerol stocks from liquid cultures as previously described. Sanger sequencing using the H1 promoter (Supplementary Fig. 1) was performed by ACGT, Inc. (Wheeling, IL, USA) to validate sequence insertion, and total plasmid sequencing was performed by the MGH CCIB DNA Core.

For consistency in plasmid production at scale, starter culture stocks were prepared as follows. Two starter cultures were inoculated with scrapings from the main glycerol stock and incubated for 12 h at 37 °C with shaking, then scaled up in 160 mL antibiotic selection media as previously described. Cultures were combined, mixed, and aliquoted into six 50 mL conical tubes (cat. no. 2231000351, Eppendorf, Hamburg, Germany), then centrifuged (1180 × g, 4 °C, 5 min) to obtain bacterial pellets. Supernatants were discarded and each pellet was resuspended in 25 mL LB broth with 100 μg/mL ampicillin and 25% glycerol. 1 mL aliquots were transferred to cryotubes and stored at -80 °C. When needed, one aliquot was thawed to RT and added to 160 mL LB broth with 100 μg/mL ampicillin and incubated 20 h at 37 °C with shaking. Plasmid DNA was extracted using the Qiagen HiSpeed Maxi Kit (cat. no. 12662, Qiagen) following the vendor’s protocol.

### Cell transfection and EV preparation

For EV preparations, cells from nitrogen storage were thawed and passaged at least twice (to a maximum of 5 times) before use. Cells cultured in T-75 flasks (cat. no. 12565349, Thermo Fisher Scientific) were grown to 70-80% confluency and transfected with 40 μg plasmid DNA using Lipofectamine 3000 (cat. no. L3000-015, Invitrogen), following the manufacturer’s protocol. After overnight incubation, the cell culture media was removed and transfected cells were washed 3 times in 1X PBS (cat. no. MT21040CV, Corning). 10 mL base medium supplemented with 10-20% vacuum-filtered (cat. no. 431162, Corning), exosome-depleted FBS (cat. no. EXO-FBS-50A-1, System Biosciences) was added to the cells, which were allowed to incubate under standard conditions for 72 h. EV-enriched cell culture media was transferred to 15 mL LoBind conical tubes (cat. no. 0030122208, Eppendorf) and centrifuged (1,000 × g, 4 °C, 10 min) in a swinging-bucket rotor to pellet residual cells and large debris. All but ∼ 1.5 mL of the supernatant was carefully transferred to new 15 mL conical tubes, then aliquoted into 1.5 mL microcentrifuge tubes (cat. no. 022431021, Eppendorf) prior to depletion of large microvesicles and other cell debris by centrifugation (10,000 × g, 4 °C, 30 min). Supernatants were consolidated into 50 mL conical tubes and mixed with 0.5 volumes of Total Exosome Isolation reagent (cat. no. 4478359, Invitrogen), then precipitated overnight at 4 °C. After mixing by inversion, the suspended precipitate was pelleted by repeated transfer of 1.5 mL into a microcentrifuge tube, centrifugation (10,000 × g, 4 °C, 5 min), and discarding of the supernatant. As a general rule, one 1.5 mL microcentrifuge tube was used for every T-75 flask used for EV enrichment. In this way, pellets were consolidated so that each pellet represented an amount of material equivalent to one T-75 flask worth of enriched media. Pellets were gently washed with 1 mL 1X PBS and then softened by overnight incubation in 100 μL 1X PBS at 4 °C. Softened pellets were resuspended by vortexing briefly, and residual precipitation reagent was removed by passing the EVs through Exosome Spin Columns (cat. no. 4484449, Invitrogen) following the manufacturer’s instructions. Samples were quantified by total protein content using a BCA protein assay (cat. no. 23228, Thermo Fisher Scientific) and diluted with 1X PBS to achieve a target dose concentration of 2 μg/ul. Dose preparations for injection were stored at 4 °C for up to 14 d.

### Identification of secreted tracer miRNA sequence

Clone 9 cells in one T-75 flask were transfected with XMc39 lentivector and EVs were isolated, as previously described. MicroRNA was extracted and cDNA was prepared, as previously described. Restriction enzyme sites and 6-nucleotide 5’ overhanging sequences were added to the tracer amplicon during PCR amplification using the following primers 5’ to 3’: XMc39 forward primer with BamHI, CCA CTT <u>GGA TCC</u> TCA CCG GGT GTA AAT CAG CTT; Universal reverse primer with EcoRI, ATC GAA <u>GAA TTC</u> GCA TAG ACC TGA ATG GCG GTA AG. Underlined sequences indicate BamHI (forward primer) and EcoRI (reverse primer) restriction enzyme sites. 20 μL reactions were prepared in triplicate in a MicroAmp Optical 96-well plate (Applied Biosystems), as follows: 2 μL cDNA, 10 μL PowerUp SYBR Green Master Mix (Applied Biosystems), 1 μL 4 μM forward primer, 0.5 μL 10 μM reverse primer, and 6.5 μL ultrapure water (cat. no. 10977-015, Invitrogen).

Amplification was performed on a QuantStudio 12K Flex using the following conditions: 50 °C for 2 min, 95 °C for 10 min, 95 °C for 15 s, 52 °C for 1 min, 40 cycles of 95 °C for 15 s and 61 °C for 1 min. All ramp rates were 1.6 °C/s. An immediate melt curve analysis was performed by heating to 95 °C with a 5 s hold every 0.3 °C step. The 3 replicate PCR reactions were pooled for a total volume of 60 μL and cleaned using a MinElute PCR Purification kit (cat. no. 28004, Qiagen) according to vendor instructions. DNA concentration of the purified eluate was quantified using a Qubit dsDNA BR assay kit (cat. no. Q32850, Thermo Fisher). Amounts of 500 ng purified tracer cDNA (insert) and 1 μg XMc39 lentivector (plasmid) were separately subjected to restriction enzyme double digestion at 37 °C overnight in 20 μL volumes containing 20 U BamHI-HF (cat. no. R3136S, New England BioLabs), 20 U EcoRI-HF (cat. no. R3101S, New England BioLabs), and ultrapure water. MinElute PCR Purification kit was used to clean the insert digest, and QIAquick PCR Purification kit (cat. no. 28104, Qiagen) was used to clean the plasmid digest according to vendor instructions. The purified digests were quantified using the Qubit dsDNA BR assay, and a ligation reaction was prepared in a 20 μL volume using 20 ng digested plasmid DNA, 2.5 μL digested insert, 800 U T4 DNA Ligase (New England BioLabs), and ultrapure water. The ligation reaction occurred for 10 min at 37 °C, then the ligase was inactivated with a 10 min incubation at 65 °C prior to chilling on ice. The ligation product was introduced into Stbl3 competent *E. Coli* cells by heat shock and plated for antibiotic selection, as previously described. Five colonies were selected and scaled up for Sanger sequencing by ACGT, Inc., as previously described.

### Electron microscopy

EVs were evaluated for morphology and contamination by the Electron Microscopy Center at Indiana University Bloomington. To prepare negative stain grid, 4 μL of sample solution was applied onto a glow-discharged 300-mesh copper grid coated with continuous carbon film (EMS, Hatfield, PA, USA). The sample solution was left for 30 s before blotted with a piece of filter paper. The grid was quickly washed using a 4-μL drop of milli-Q (MilliporeSigma) water and stained with 4 μL of negative stain solution composed of either 1% (w/v) uranyl acetate (EMS) with 0.5% (w/v) trehalose (MilliporeSigma) or 1% (w/v) ammonium molybdate (MilliporeSigma) with 0.5% (w/v) trehalose. The excess stain solution was removed by filter paper and the grid was put aside to allow air dried. Grids were imaged on a 120-kV JEM-1400Plus (JEOL USA Inc., Peabody,MA, USA) transmission electron microscope equipped with 4k × 4k OneView camera (Gatan Inc., Pleasanton, CA, USA).

### Nanoparticle tracking analysis

EV preparations were analyzed for size distribution with dynamic light scattering using the Particle Metrix ZetaView platform (Particle Metrix, Meerbusch, Germany). Data acquisition was performed at RT using dilutions of EVs in 1X PBS. Using EV preparations diluted to a protein content of 2 μg/μL, a starting dilution of 15 μL in 1 mL of PBS was used and then further diluted to achieve empirical particle concentrations within the acceptable range of the analysis software. Nanoparticle tracking analysis measurements were recorded and analyzed at 11 positions per sample with the ZetaView analysis software (Particle Metrix). Size distributions were obtained from 3 biological replicates (EVs prepared on 3 separate occasions).

### Gel electrophoresis and western blot analysis

Following collection of EV-enriched media, adherent cells were washed in triplicate using 1X PBS then detached by incubation in trypsin (cat. no. MT25053CI, Corning) for about 5 min at RT. Detached cells were collected with complete growth medium to inactivate trypsin, then pelleted by centrifugation at 200 × g for 5 min at 4 °C. Cell pellets were washed in triplicate by resuspension in PBS and repeat centrifugation. Aliquots from the final resuspension were used to count cells using a Fuchs Rosenthal hemocytometer (cat. no. DHC-F01, Incyto, Republic of Korea). Final cell pellets were lysed on ice for 10 min in 1X RIPA buffer (cat. no. 9806S, Cell Signaling Technology, Danvers, MA, USA) with added protease inhibitors (cat. no. 78429, Thermo Fisher Scientific). Following centrifugation at 14,000 × g for 10 min at 4 °C, lysate supernatants were collected and quantified by BCA assay.

50 μg aliquots of both EVs and whole cell lysates (WCL) were stored at -20 °C until analyzed. Stored aliquots were thawed in LDS sample buffer (cat. no. NP0007, Invitrogen) with NuPAGE sample reducing agent (cat. no. NP0004, Invitrogen), then heated at 75 °C for 10 min. For probing the tetraspanins, additional aliquots were prepared without the addition of reducing agent. Denatured samples, along with Precision Plus Protein Kaleidoscope Prestained Protein Standards (cat. no. 1610375, Bio-Rad, Hercules, CA, USA) and MagicMark XP Western Protein Standards (cat. no. LC5602, Invitrogen), were resolved on precast NuPAGE 4-12% Bis-Tris midi 12+2-well protein gels (cat. no. WG1401BOX, Invitrogen) at 200 V for 40 min in NuPAGE 2-(*N*-morpholino)ethanesulfonic acid (MES) running buffer (cat. no. NP0002, Invitrogen), supplemented with NuPAGE antioxidant (cat. no. NP0005, Invitrogen) in the case of reduced samples. For western blot analysis, gels were transferred to 0.45 μm polyvinylidene difluoride membranes (cat. no. IPFL00010, MilliporeSigma) using the Criterion blotter and Towbin buffer (cat. no. 1704070 and 1610771, Bio-Rad) at 10 V overnight in a cold room, with stirring. Protein transfer was verified using Ponceau S staining (cat. no. P3504, MilliporeSigma) of the membranes. Membranes were destained, then blocked in 3% BSA/TBS-T (TBS containing 0.1% Tween 20; cat. nos. BP9706100, BP2471500, BP337100, Thermo Fisher Scientific) for 45 min at RT, with rocking. Where applicable, membranes were cut into strips using visible molecular weight markers as guides. Membranes or membrane strips were probed overnight at 4 °C using mouse monoclonal primary antibodies diluted in 1% BSA/TBS-T, with rocking. Membranes were washed for 5 min in TBS-T, in triplicate, then incubated with anti-mouse IgG horseradish peroxidase(HRP)-linked secondary antibodies (cat. no. 7076, Cell Signaling Technology) diluted 1:3,000-1:10,000 in 5% non-fat milk/TBS-T for 2 hours at RT. Membranes were washed thrice in TBS-T as before, then incubated in SuperSignal West Femto Maximum Sensitivity Substrate (cat. no. 34094, Thermo Fisher Scientific) for 5 min. Membranes were placed in-between layers of a page protector and imaged using a ChemiDoc MP Imaging System (Bio-Rad).

Mouse monoclonal primary antibodies included: anti-CD63 (clone MX-49.129.5, 2 μg/mL, cat. no. sc-5275, Santa Cruz Biotechnology, Dallas, TX, USA), anti-CD81 (clone 1.3.3.22, 2 μg/mL, cat. no. MA5-13548, Invitrogen), anti-tsg 101 (clone C-2, 2 μg/mL, cat. no. sc-7964, Santa Cruz Biotechnology), anti-Alix (clone 3A9, 2 μg/mL, cat. no. MA1-83977, Invitrogen), anti-ApoA-I (clone 069-01, 1 μg/mL, cat. no. sc-58230, Santa Cruz Biotechnology), anti-Histone cluster 1 H3D (clone 6H8, 1 μg/mL, cat. no. sc-134355, Santa Cruz Biotechnology), anti-cytochrome c1 (clone A-5, 2 μg/mL, cat. no. sc-514435, Santa Cruz Biotechnology), anti-GM130 (clone B-10, 2 μg/mL, cat. no. sc-55591, Santa Cruz Biotechnology), anti α-actinin (clone H-2, 2 μg/mL, cat. no. sc-17829, Santa Cruz Biotechnology), anti-eIF2C (Argonaute1-4; clone B-3, 2 μg/mL, cat. no. sc-376696, Santa Cruz Biotechnology), anti-hnRNP A2/B1 (clone b-7, 2 μg/mL, cat. no. sc-374053, Santa Cruz Biotechnology).

### Lipidomic mass spectrometry

The MRM-profiling methodology was used as previously described for the lipidomic analysis of bovine oocytes and preimplantation embryos, and for evaluating aging of fat tissue^13,14^. The experiments were performed using an Agilent 6410 QQQ mass spectrometer (Agilent Technologies, Santa Clara, CA, USA) with a micro-autosampler (G1377A). A lipid extraction was performed on the EV samples using the Bligh & Dyer protocol^15^. This protocol called for a volume of 200 μL of buffer containing the EV protein combined with 450 μL and 250 μL of methanol and chloroform, respectively, in a microtube. After allowing the samples to sit at RT for 15 min, 250 μL of ultrapure water and chloroform were added. They were then centrifuged in order to amplify the separation of the lipid, metabolite, and protein phases based on differences in polarity. The lipid (bottom) layer was extracted and dried under a stream of nitrogen and stored at -80 °C until MS analysis.

The dried samples were then resuspended in an appropriate volume of acetonitrile (ACN)/methanol/ammonium acetate 300 mM, v/v/v, 6.65:3:0.35 (injection solvent). The volume of 8 μL of EV diluted lipid extract was injected into the electrospray ionization (ESI) source of the MS. The capillary pump connected to the autosampler operated at a flow rate of 10 μL/min and a pressure of 100 bar. Capillary voltage on the instrument was 3.5-5 kV and the gas flow was 5.1 L/min at 300 °C.

MRM-profiling is a two-phase process containing both a discovery and screening phase. In each phase, MRMs are used to investigate the functional groups of a sample based on neutral loss and precursor MS/MS scans. The representative sample pool used in the discovery phase consisted of 14 different EV samples from rat cell lines. For this phase, using methods previously reported by de Lima et al., we applied a list of 1,419 MRMs from 10 lipid classes: phosphatidylcholine (PC)/sphingomyelin (SM), phosphatidylethanolamine (PE), phosphatidylinositol (PI), phosphatidylglycerol (PG), phosphatidylserine (PS), ceramide, cholesteryl ester (CE), acyl-carnitine, free fatty acid (FFA), and triacylglycerol (TAG)^13^. The monitoring of these classes was based on precursor ions of lipids listed in the Lipid Maps Database (http://www.lipidmaps.org/) and product ions common to each given lipid class.

The raw MS data, MRM transitions and intensities, were processed using in-house scripts in order to generate a list of MRM transitions and their respective ion intensities. Comparison of the absolute ion intensities for the EVs to a blank sample (injection solvent) was then assessed and the MRMs that depicted an ion intensity at least 30% higher than the blank were selected. The top 200 MRMs were selected to be used in the screening phase and these were monitored over a period of 2 min per sample. The screening method included MRMs from five lipid classes (PC and SM, Cholesteryl esters, ceramides, PE) and a single metabolite (acyl-carnitine) class.

### RNA Extraction and cDNA synthesis

XMIR-NT positive control RNA oligonucleotide was purchased from SBI. XMc39 positive control RNA oligonucleotide was purchased from IDT: 100 nmole, UCA CCG GGU GUA AAU CAG CUU GCC UAG GAG GAG. RNA extractions were performed using the Qiagen miRNeasy Mini Kit (cat. no. 217004, Qiagen), and 1.5 mL DNA LoBind microcentrifuge tubes (cat. no. 022431021, Eppendorf) were used when possible. Frozen aliquots of dose preparations and clarified plasma were thawed to RT in batches, so that miRNA was extracted from animal-matched samples in parallel. Since it is impractical to quantify miRNA extracted from small volumes of plasma, we adapted a volume normalization method used in one of our previous studies^16,17^. At the time of freezing, samples were apportioned as 50 μL volumes in 1.5 mL microcentrifuge tubes. Upon removal from -80 °C storage, 250 μL Qiazol reagent was added to each sample. After thawing to RT in the presence of Qiazol, samples were vortexed at maximum speed for 5 s, then incubated at RT for 5 min. After addition of 50 μL chloroform (cat. no. 319988, Sigma-Aldrich), samples were vortexed for 12 s and incubated at RT for 3 min. Phase separation was performed by centrifugation at 12,000 × g for 15 min at 4 °C. For each sample, 150 μL of the upper aqueous phase was transferred to a new microcentrifuge tube and mixed with 225 μL 100% ethanol (cat. no. BP2818500, Fisher Scientific) by gently pipetting 4X. Ethanolic mixtures were transferred to RNA-binding columns and washed with Buffers RWT and RPE according to vendor instructions, including the optional high-speed centrifugation. Elution was performed with 30 μL ultrapure water pre-heated to 60 °C, incubation at RT for 5 min, then centrifugation at 8,000 × g for 1 min. A second elution was performed in the same manner, but with a 5 min final centrifugation. The final eluate volume was 60 μL.

First strand cDNA was prepared using the qScript miRNA cDNA Synthesis kit (cat. no. 95091-025, Quantabio, Beverly, MA, USA). DNA LoBind products were used for all steps, including 1.5 mL microcentrifuge tubes, 0.5 mL microcentrifuge tubes (cat. no. 2231000341, Eppendorf), and 96-well plates (cat. no. 2231000368, Eppendorf). Steps were performed according to the vendor’s protocol, with the following specifics. 7 μL volumes of RNA were added to the wells of a 96-well plate. An appropriate amount of poly(A) polymerase was added to poly(A) tailing buffer in a microcentrifuge tube, vortexed and centrifuged briefly to settle, then added to samples. Sample wells were capped (cat. no. 0030124839, Eppendorf) and the 96-well plate was vortexed to mix and centrifuged to settle. Poly(A) tailing reactions were performed on a Veriti thermal cycler (Applied Biosystems) according to the vendor’s protocol using the 20 min option for the 37 °C incubation. Reverse transcriptase and cDNA reaction mix were prepared and added to the poly(A) tailing reactions in a similar manner. After the reverse transcription (RT) reaction, samples were held at 4 °C until ddPCR, then stored at -80 °C.

### Droplet digital PCR

Droplet digital PCR (ddPCR) was performed using the QX200 AutoDG ddPCR system (cat. no. 1864100, Bio-Rad) and ddPCR Supermix for EvaGreen (cat. no. 1864034, Bio-Rad) according to the vendor’s protocol. A forward primer of unknown sequence for XMIR-NT was initially purchased from System Bioscience (Palo Alto, CA, USA), then later custom ordered from IDT after plasmid sequencing. Primer sequences were as follows: XMIR-NT forward primer, GAG GGC GAC TTA ACC TTA G. XMc39 forward primer, TCA CCG GGT GTA AAT CAG C; Universal reverse primer, GCA TAG ACC TGA ATG GCG GTA.

Preamplification was performed using ddPCR Supermix for EvaGreen as follows. Amplification reactions of 15 μL were prepared in an Eppendorf DNA LoBind 96-well plate. Each reaction consisted of 1.5 μL cDNA, 7.5 ul ddPCR Supermix for EvaGreen, 4.5 μL ultrapure water, and 1.5 μL 2.5 µM forward primer. Undiluted cDNA was used from the RT reaction. The final forward primer concentration was 250 nM. No reverse primer was added, as oligo-d(T) carryover from the RT was sufficient to act as a reverse primer. Samples were amplified in a Veriti thermal cycler using the following conditions: 95 °C for 5 min, 5 cycles of 95 °C for 30 s and 58 °C for 60 s (100% ramp rate), 4 °C for 5 min, 90 °C for 5 min, and hold at 4 °C. Since Taq polymerase is activated during the preamplification, samples were prepared for ddPCR as soon as the temperature of the samples reached 4 °C.

We prepared ddPCR reactions according to Bio-Rad’s specifications, with some modifications. Briefly, 2.5 μL of each preamplification reaction was added to a ddPCR reaction with a final volume of 25 μL and forward and reverse primer concentrations of 200 nM each. First, a master mix of ddPCR Supermix and primers was prepared and aliquoted into a LoBind 96-well plate (the “supermix plate”) held at 4 °C on a cold block. Next, the preamplification plate was placed on a 4 °C cold block and samples were transferred to the wells of the supermix plate. The supermix plate was briefly vortexed and centrifuged to mix and settle contents. Droplets were prepared using 20 μL of each supermix sample.

Droplets were created using the automated droplet generator or the manual QX200 droplet generator (cat. no. 1864002, Bio-Rad), depending on availability and number of samples. Automated droplet generation oil for EvaGreen (cat. no. 1864112, Bio-Rad), DG32 automated droplet generator cartridges (cat. no. 1864109, Bio-Rad), QX200 droplet generation oil for EvaGreen (cat. no. 1864005, Bio-Rad), DG8 droplet generator cartridges (cat. no. 1864008, Bio-Rad), DG8 gaskets (cat. no. 1863009), and ddPCR 96-well plates (cat. no. 12001925, Bio-Rad) were used as needed. Droplets were transferred to a ddPCR 96-well plate and heat sealed with foil (cat. no. 1814040, Bio-Rad) using a PX1 plate sealer (cat. no. 1814000, Bio-Rad).

Droplets were amplified to endpoint using the following cycling conditions on a C1000 Touch thermal cycler (cat. no. 1851197, Bio-Rad): 95 °C for 5 min, 40 cycles of 95 °C for 30 s and 56 °C for 60 s (default ramp rate; 2.5 °C/s), 4 °C for 5 min, 90 °C for 5 min, and hold at 4 °C. Following thermal cycling, droplets were scanned using the QX200 Droplet Reader (Bio-Rad). Analysis was performed using QuantaSoft Analysis Pro software (Bio-Rad).

### *In vivo* kinetic experiments

EVs from transfected clone 9 cells expressing XMc39 tracer were used to establish a target dose amount for animal experiments. EVs were prepared in bulk and quantified by protein concentration as previously described. RNA extracted from 100 μg EVs (2 μg/μL) was diluted 1:100 in water and used to prepare cDNA for ddPCR analysis, as previously described. Using the approximate total blood volume of a 300-450 g male Sprague Dawley rat^18^, we determined that 1,000 μg EVs would result in an initial concentration (C_0_) near the upper limit of ddPCR detection. A preliminary *in vivo* time course was performed to validate the calculated dose amount, and to establish experimental duration.

High and low standards were produced to capture the batch variability of RNA extraction and analysis, as follows. Citrated blood from two exsanguinated naïve animals was pooled. “Positive” plasma was prepared by mixing 1,333 μg clone 9 EVs with 11.5 mL blood by gentle inversion. “Negative” plasma was prepared in parallel using an identical volume of 1X PBS in place of EVs, but otherwise using the same protocol. Blood was transferred to multiple 1.5 mL microcentrifuge tubes (Eppendorf) and plasma was separated by centrifugation (2,000 × g, 20 min, 4 °C) in a Microfuge 20R (cat. no. B30149, Beckman Coulter, Brea, CA, USA). Plasma was transferred to new 1.5 mL microcentrifuge tubes and further clarified by centrifugation (10,000 × g, 10 min, 4 °C). “Positive” and “negative” plasma were pooled in 15 mL conical tubes (cat. no. 2231000349, Eppendorf), respectively. The “high” standard was developed by diluting “positive” plasma with “negative” plasma and empirically determining the concentrations by ddPCR until a desirably high signal was achieved. The “low” standard was prepared by diluting the “high” standard 30-fold with “negative” plasma. “High” and “low” standards were evaluated by ddPCR to ensure they fell within the expected dynamic range of our kinetic experiments, i.e. not to exceed the highest and lowest expected observations. 50 μL aliquots of high and low standards were stored in 1.5 mL low-binding microcentrifuge tubes (Eppendorf) at -80 °C.

For replicate experiments, catheter access was achieved by placing conscious animals in a rodent restrainer and using 1 mL tuberculin slip-tip syringes (cat. no. 309626, BD) with attached blunt 22 ga dispensing needles (cat. no. JG220, Jensen Global, Santa Barbara, CA, USA). Animals were weighed prior to dose administration and then again at euthanasia.

For blood collection, each syringe was carefully prepared under sterile conditions: The included needle was discarded, the plunger was drawn back to 0.1 cc, and 20 µL 4% sodium citrate was pipetted into the syringe’s slip-tip prior to placement of an autoclaved blunt dispensing needle. At the time of collection, 200 μL blood was collected by retracting the plunger to 0.3 cc. The syringe was disconnected from the catheter, plunger retracted to 0.5 cc, and citrated blood mixed by 2-3 gentle inversions. The catheter was flushed with 0.25 cc saline and locked with 0.1 cc 4% sodium citrate. Blood samples were transferred to a 0.5 mL low-binding microcentrifuge tubes (cat. no. 2231000341, Eppendorf) as soon as possible after locking the catheter.

For dosing, the target dose amount for all experiments was 1,000 μg protein equivalent in a volume of 500 μL 1X PBS. Twenty percent excess was prepared to account for the dead space of the luer lock and allow for potential loss of fluid while clearing the syringe of air bubbles. At the time of dosing, lock solution and 0.1 mL blood were drawn from the catheter into a syringe and discarded. A negative control blood sample was collected immediately prior to injecting 500 μL EV dose. A dose aliquot of 50 μL was reserved from the remaining fluid and stored at -80 °C for analysis and data normalization. The catheter was flushed with 500 μL bacteriostatic saline and locked with 100 μL 4% sodium citrate.

Blood samples were collected from each animal at 2, 7.5, 15, 30, 60, 120, 240, 480, 960, and 1440 min after dosing. Each collection involved the following steps: discarding of lock solution and 0.1 mL blood, collection of 0.2 mL blood, pulsatile flushing of the catheter with 0.25 mL bacteriostatic saline, and locking of the catheter with 0.1 mL 4% sodium citrate. Blood plasma was separated from the blood (2,000 × g, 20 min, 4 °C) and then clarified (10,000 × g, 10 min, 4 °C) by centrifugation. Two 50 μL aliquots were transferred to 1.5 mL low-binding microcentrifuge tubes and stored at -80 °C for downstream analysis.

### Tracer miRNA time course stability assay

Immediately prior to conducting an *in vivo* kinetic time course, detailed above, blood was collected from one male Sprague Dawley rat by cardiac puncture. Briefly, the animal was euthanized by isoflurane inhalation (5% induction, 5% maintenance). A laparotomy was performed, followed by a bilateral anterolateral thoracotomy. One 20 mL syringe (cat. no. 309661, BD) pre-filled with 1 mL 4% sodium citrate (Fenwal) was used to obtain 10 mL blood from the exposed heart. The citrated blood was mixed by gentle inversion and 8 mL was transferred to a 15 mL LoBind conical tube (Eppendorf), then placed in a heated water bath set to 37 °C. The *in vitro* and *in vivo* time course experiments were performed in parallel, beginning with the administration of EVs. Immediately after dosing the conscious rat for the *in vivo* time course, 150 μL (300 μg) of the same EV dose preparation was spiked into the *in vitro* anticoagulated blood and mixed by gentle inversion. Immediately subsequent to each blood collection from the conscious rat, 220 μL of blood was collected from the conical tube; since the *in vitro* blood was previously mixed with citrate, 220 μL was collected to account for both the 200 μL sampled from the conscious animal and 20 μL sodium citrate added to these sampling syringes, as detailed previously. After each blood collection, the citrated blood in the conical tube was mixed by gentle inversion to prevent settling of the red blood cells over time. *In vitro* blood samples were collected at the same time points, up to 240 min, and handled in the same way alongside the *in vivo* blood samples all the way through ddPCR analysis.

### Preparation of standard curve

Two standard curves were independently performed using serial dilutions of the high standard. Since the high standard was designed to achieve a maximum copy concentration in an intermediate-high range (∼25,000 copies/20 μL), a 5-fold concentration was prepared by performing Qiazol phase separation for each of 5 aliquots and binding the RNA precipitates to the same silica membrane column prior to elution. From this 5X high standard, twofold dilutions were prepared using miRNA from naïve rat plasma as the diluent. Concentrations were obtained by ddPCR.

### Data normalization

To account for technical variability, high and low standards were included with every set of samples analyzed by ddPCR. We normalized the data using our standard curve as follows. Standard curve copy numbers were plotted against their concentration factor; the concentration factors ranged from ∼0.001 to 5. A linear regression of the standard curves was performed with Excel 2019 (Microsoft Corporation, Redmond, WA, USA), and reference standard copy numbers were calculated for concentration factors of 1, corresponding to the 1X high standard, and 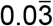 corresponding to the low standard which is a 30-fold dilution of the high standard. For every set of samples, the internal high and low standards were used to normalize observed copy numbers to the reference standards. Copy number concentrations were then converted to EV protein concentrations and normalized against the dose aliquot for each set of samples, taking all dilution factors into account (Supplementary Fig. 5). Normalizing copy numbers to EV protein concentrations effectively accounts for differences between cell lines and potential variability between EV preparations.

### EV pharmacokinetic modeling

Modeling EV disposition following IV administration was performed using a population pharmacokinetic approach. Phoenix 64 build 8.1.0.3530 (Certara, Princeton, NJ, USA) was used to support non-linear mixed effects analysis with first order conditional estimation - extended least squares (FOCE ELS) to estimate population-level parameters with associated inter-animal variability on those parameters. Initial parameter estimates were made using the “initial estimates” function in Phoenix to manually create the best fit lines to the observed data. Subsequently, each sequence of parameter estimation was limited to a maximum of 1,000 iterations. Observed concentrations were fit to the exponential form of equations describing two-compartment and three-compartment model structures (Fig. 4b). Equations were parameterized according to clearance between compartments and the compartment volumes. Inter-individual (IIV) random effects for the various structural parameters were included as a diagonal matrix initially. These random effects are reported as percent variance from a log-normal distribution of individual subject parameter estimates, the basis of which is the exponential relationship, P_i_ = P_tv_ × exp(η_i_), where P_i_ is the parameter estimate for the ith individual, P_tv_ is the population typical value, and η_i_ (eta) is the deviation from the population value for the ith subject. Correlation of IIVs among parameters was evaluated graphically to support the need to estimate covariance of random effects between parameters. A multiplicative (proportional) residual error model was applied using the relationship, Cobs = C * (1 + CEps)), where Cobs is the observed concentration, C the model predicted concentration and CEps the difference between Cobs and C. Covariates were multiplied to population-parameter estimates (thetas) exponentially as theta *e^covariate^. Evaluation of the final 3 compartment model with cell line covariates consisted of a prediction-corrected visual predictive check (pcVPC) of 1,000 simulations based on the final parameter estimates. Bootstrap analysis was used to evaluate parameter stability. For the pcVPC, a log-additive residual error model was used in place of a multiplicative error model. The log-additive model is the same as a multiplicative model, except that it prevents simulations resulting in negative EV concentrations, as negative concentrations are not possible. Simulated concentrations from the pcVPC were stratified by cell line, and the concentrations binned by k-means (the mean of the times). Median and associated 5% and 95% confidence limits of the observed EV concentrations were superimposed with their corresponding median predicted values and associated 5-95% intervals of these median predictions. The bootstrap analysis consisted of 1,000 samples with replacement from the original set of animals (each sample containing the same number of animals as the original study).

### Statistics

Lipidomic analysis was performed with MetaboAnalyst 4.0 (www.metaboanalyst.ca) using the following options. Sample ion counts were normalized by sum and auto-scaled. One-way ANOVA and Fisher’s LSD post-hoc analysis were performed to select top scoring lipids (unadjusted *P* < 0.05) for PCA and heat map Sample sizes for this study were determined using data from Morshita et. al^4^. A sample size of 10 rats would have 80% power to detect a 30% change in exosome clearance using an unpaired *t*-test and 5% type 1 error rate. This is estimated based on the calculation of EV clearance to be 0.52 ml/min, and a conservative estimate of 25% variability (given the limited data available). EV clearance was calculated using the following equations: 100% ID = 37 kBq; 37 kBq/100% ID × 3.2 (%ID × hr / mL) = 1.184 kBq × hr / mL = AUC; CL = d / AUC; CL = 37 kBq / 1.184 kBq × hr /mL; CL = 31.25 mL / hr; thus CL = 0.52 mL / min. CL = clearance, d = dose, hr = hour, AUC = area under the concentration-time curve; kBq = kilo Becquerel. We also determined this sample size and sampling frequency per animal was adequate to support non-linear mixed effects analysis.

Elimination half-life (T ½), compartment distribution half-life, and AUC were determined from the Phoenix post-hoc data for the final model. Elimination T ½ for each sample ln 2/(Cl/(V + V2 + V3)), compt 2 distrib T ½ = ln2 / (Cl2/V2). Compt 3 distrib T ½ = ln2 /(Cl3/V3). JMP Pro 14 was used for statistical analysis. Given a sample size of 10 and without the assumption of normal distribution or equal variance, Wilcoxon and Kruskal-Wallis rank-sum tests were applied as a conservative non-parametric approach to determining significant differences between cell lines (P < 0.05). If significance was met by the Wilcoxon/Kruskal-Wallis test, then the Steel-Dwass method was applied to evaluate for significant differences between cell lines. Steel-Dwass makes non-parametric comparisons for all pairs and takes into account multiple comparisons similar to Tukey’s Method for parametric data.

## Results

### Preparation of labeled extracellular vesicles

In order to discriminate exogenously administered EVs from endogenous background in rats, we incorporated a tracer miRNA sequence that did not share homology with known rat miRNAs. The chosen tracer miRNA was expressed using a commercial lentivector which appends an exosome localization motif^19^ to the resulting mature miRNA. For early optimization experiments, we used a prepackaged, proprietary non-targeting sequence from the vendor designated “XMIR-NT”. During development, however, we encountered constraints that required a known sequence. We selected *C. elegans* miR-39-3p (cel-miR-39) for cloning into the same lentivector (Supplementary Fig. 1), designated “XMc39”. Because of its non-homology with certain species, cel-miR-39 is commonly used as a quality control spike-in for miRNA PCR experiments involving biofluids from humans, rats, and other mammals^17,20,21^. The validated XMc39 plasmid was transfected into 3 established rat-derived cell lines (clone 9 liver hepatocyte, RFL-6 lung fibroblast, and RMC kidney mesangial cells). The transfected cells were cultured in EV-depleted cell culture medium for several days in order to enrich the media with secreted vesicles. EVs were isolated from enriched media using a commercial chemical isolation reagent (Fig. 1). Compared to ultracentrifugation, chemical reagents allow for substantially greater yield when retrieving EVs from biofluids and cell culture supernatants at the expense of purity^22^. Co-precipitation of medium to large vesicles^3^ was minimized by modifying the manufacturer’s protocol with an additional clarification step^23,24^ prior to addition of reagent. Remaining chemical reagent which can cause vesicle aggregation (Fig. 2a) was removed by careful washing of the pellet, resuspension, and filtration through low molecular weight size exclusion columns. Nanoparticle tracking analysis and transmission electron microscopy confirm that 86.5% ± 1.5% (mean ± S.E.) of all EVs are in the 45-195 nm size range (Figs. 2a,c; Supplementary Fig. 2), which is typical of exosome-enriched small EV (sEV) preparations, and are deprived of aggregates (Fig. 2a). Western blot analysis (Fig. 2b) probed for the appropriate presence and absence of various sEV-associated proteins and relevant co-precipitated non-sEV contaminants^3,19^ in comparison to total lysates from their originating cell lines, based on MISEV 2018 recommendations^3^. The tetraspanins CD63 and CD81 (category 1a^3^) were represented in all EVs, consistent with other reports of reagent-based EV isolation methods^23,25-29^. Cytosolic membrane-binding proteins Alix and tsg 101 (category 2a^3^) were detectable in all EV samples. The apolipoprotein and mitochondrial markers ApoA-I (category 3a^3^) and cytochrome c (category 4b^3^) were present in EV samples, though not particularly abundant relative to originating cell lines. The secretory pathway (Golgi) marker GM130 (category 4c^3^) was absent in EV samples. Histone H3.1 (category 4a^3^) was particularly enriched in two EV samples. It has been reported that histones are associated with sEVs^30-32^, but on this point there is recent disagreement^33^. The cytoskeletal marker α-actinin (category 4d^3^) was present in all samples, indicating the possible co-precipitation of autophagosomes. Relevant to the nature of our tracer, two secreted non-vesicular miRNA-binding proteins (category 5^3^) were also assayed. Argonaute1-4 were detectable in two EV samples. Non-sumoylated hnRNP A2/B1 (∼35 kDa), a specific miRNA-binding protein implicated in the mechanism by which our tracer miRNA is selectively loaded into EVs^19^ (Supplementary Fig. 1), was barely detectable in one EV sample, whereas a higher molecular weight band was visible in all EV samples at higher exposures (Supplementary Fig. 21b). Mass spectrometry confirmed that our EV preparations were enriched in sphingolipids and cholesterols (Supplementary Fig. 3, Supplementary Data 1), a hallmark of exosomes^34^, and indicated differences in lipid composition between EVs from each cell line (Fig. 2d).

**Figure 1:**
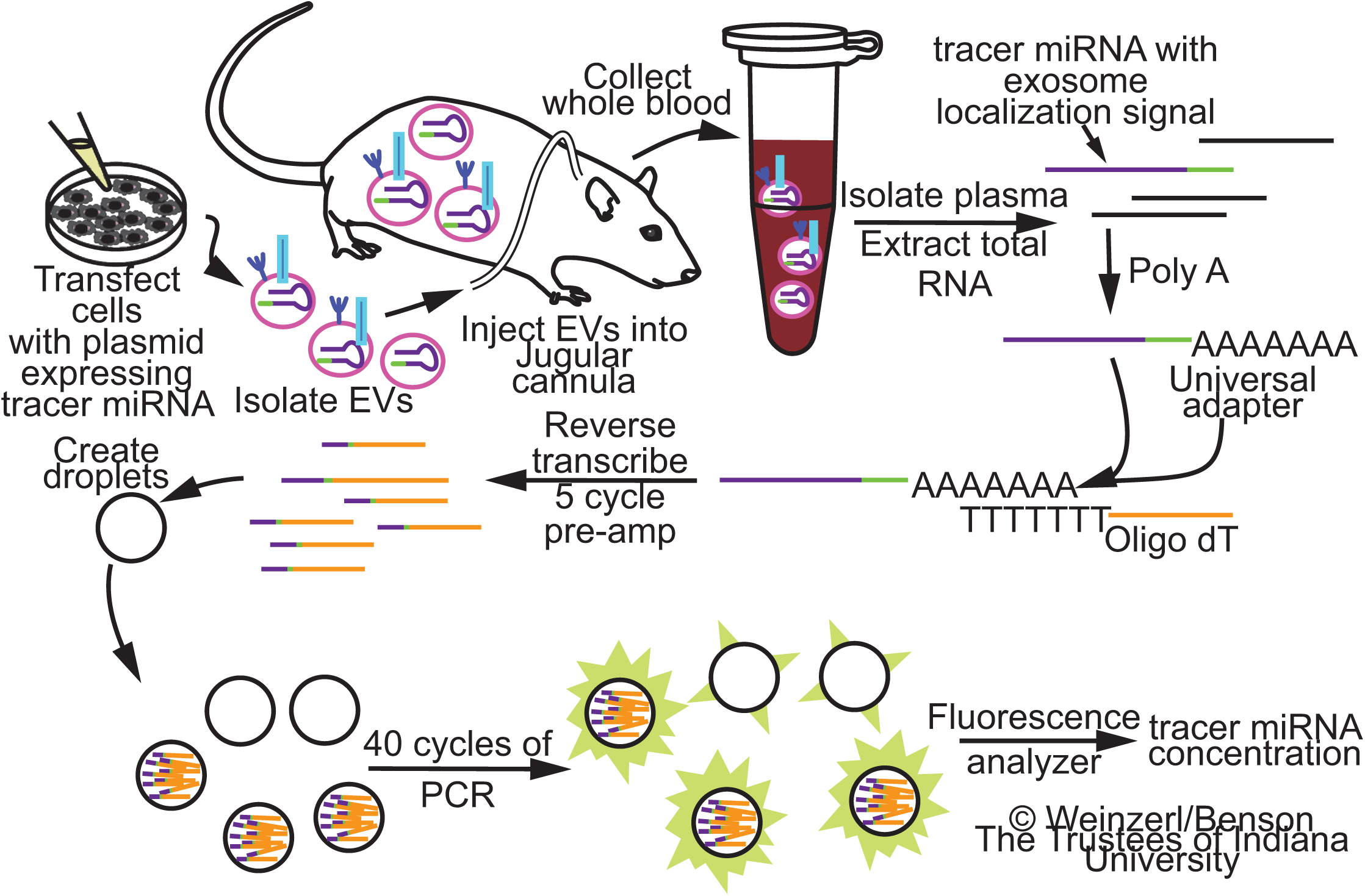
Overview of the workflow. Cultured cells are transfected with an expression vector encoding a non-homologous tracer miRNA stem-loop sequence with an exosome localization signal. EVs are isolated from the enriched media and administered to a conscious rat through a jugular cannula. Blood samples are collected at various time intervals, and total miRNA is extracted from the plasma. Complementary DNA is synthesized by polyadenylation and priming of the reverse transcriptase with an oligo(dT) adapter. 5-cycle PCR pre-amplification and Droplet Digital PCR are performed using primers directed to the tracer miRNA and oligo(dT) adapter. Tracer miRNA concentration is quantified using an EvaGreen fluorescence-based assay.

**Figure 2:**
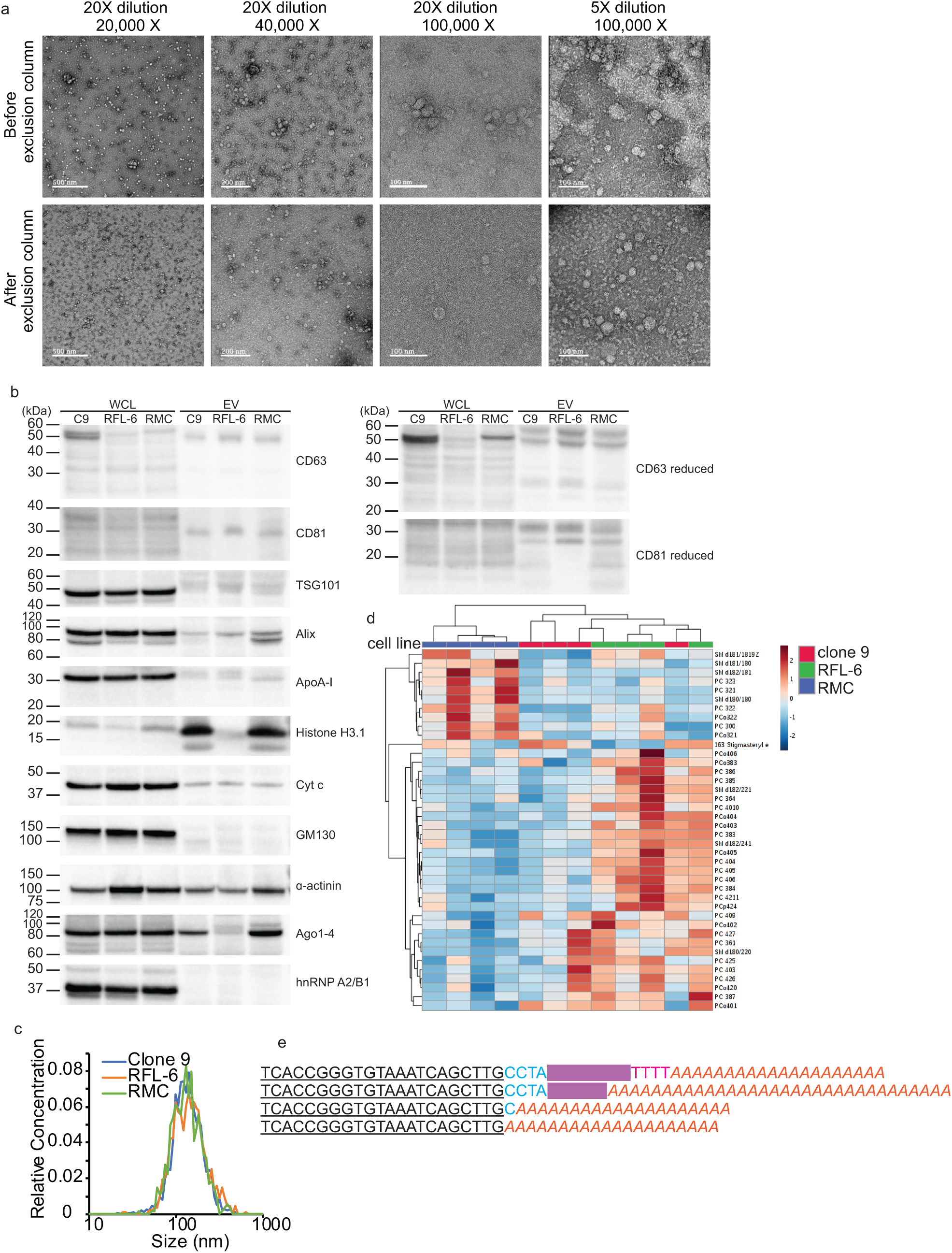
Characterization of EVs. **a**, Transmission electron micrographs of EVs before and after purification by size exclusion centrifugation. The first three columns represent different imaging magnifications (20,000X, 40,000X, and 100,000X) with a sample dilution of 20X; the fourth column represents 100,000X magnification with a sample dilution of 5X. The top row represents unpurified samples, and the bottom row represents purified samples. **b**, Western blots for EV and non-EV markers in whole cell lysates (WCL) and EV preparations (Clone 9, RMC, RFL-6). Molecular weight markers are designated by lines on the left of each blot. **c**, Average size distributions of EVs from cultured clone 9 hepatocytes, RFL-6 lung fibroblasts, and RMC mesangial kidney cells. Average distribution was produced by taking average bin counts of 3 replicates per cell line. **d**, Heat map representing the top 31 EV lipids, clustered by cell type (n = 4 for each cell line). Grouping was performed by unsupervised hierarchical cluster analysis (Euclidean distance, Ward linkage) of ion counts normalized to sum and auto-scaled. **e**, EV-associated tracer miRNA products with variable length 3’ sequences. An expression vector-specific sequence (blue) separates the mature cel-miR-39-3p sequence (bolded and underlined) from the exosome localization signal (redacted in magenta). The first sequence includes a partial poly-T transcription termination sequence (red), which is encoded in the expression vector. Long poly-A tails (orange italics) are added during miRNA cDNA synthesis.

**Figure 3:**
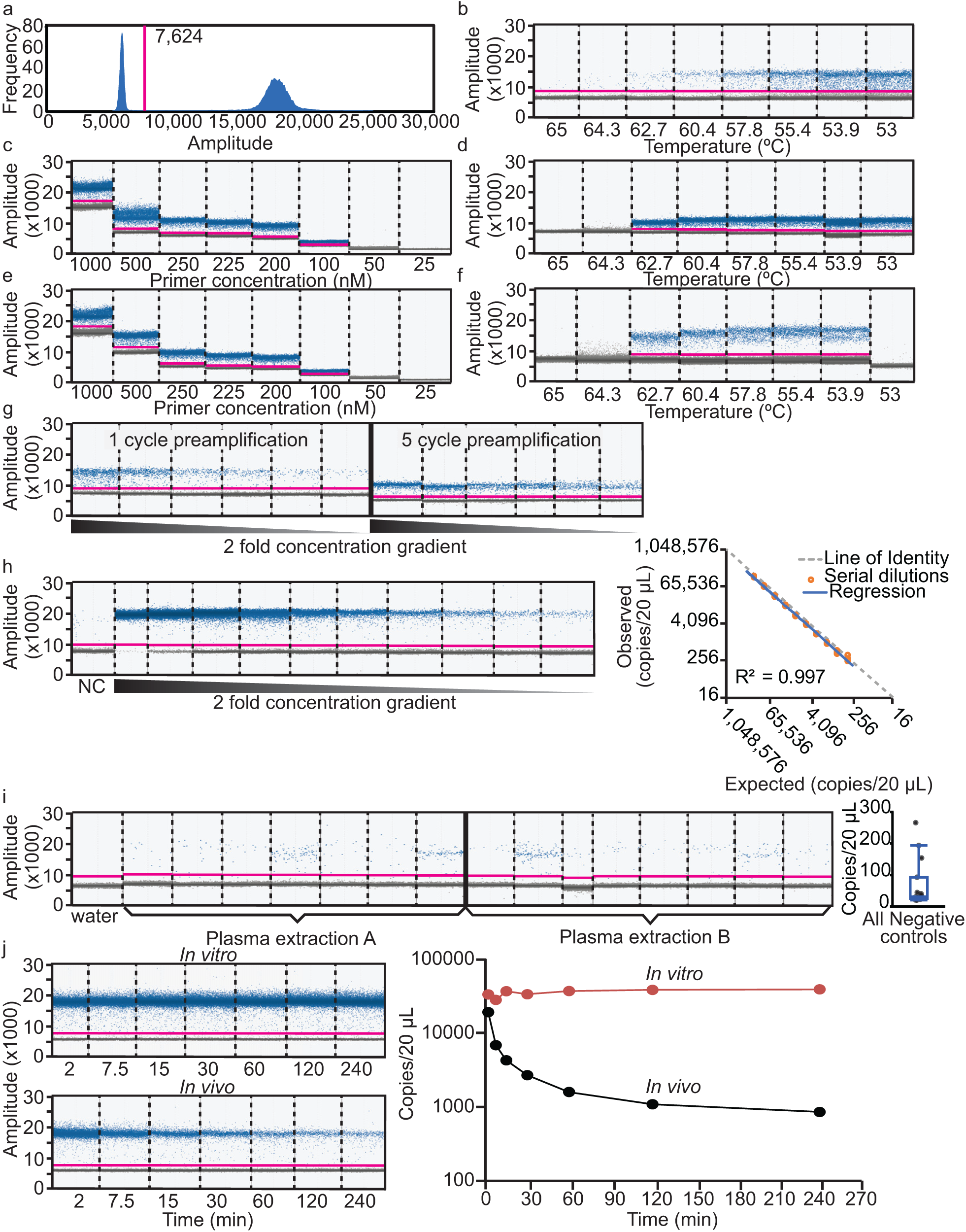
ddPCR assay design and optimization. **a**, Representative histogram of EvaGreen fluorescence used to set a threshold (magenta line) between positive (left peak) and negative (right peak) droplets. **b**, Amplitude scatterplot of initial annealing temperature (Ta) gradient (65 – 53 °C) using XMIR-NT primer (100 nM) and fixed amount of XMIR-NT cDNA template. **c**, Primer gradient (25 - 1000 nM); Ta = 60 °C. **d**, Ta gradient (65 – 53 °C) using 250 nM primer; negative control (water) in the 65 °C well. **e**, Primer gradient; Ta = 58 °C. **f**, XMc39 (200 nM) Ta gradient (65 – 53 °C). **g**, Comparison between 1- and 5-cycle PCR pre-amplification. **h**, Standard curve and evaluation of assay linearity. Concentrations from two-fold serial dilutions of XMc39 RNA (left) were used to plot expected vs. observed values (right). Individual observed concentrations (orange circles) and linear regression (blue line) closely aligned with the line of identity (dashed gray line). **i**, Technical replication of negative control plasma samples; ddPCR (left) using n = 7 and n = 8 technical replicates from two separate RNA extractions, and boxplot (right) of concentrations. **j**, Evaluation of EV-encapsulated tracer miRNA stability over time in anticoagulated whole blood at 37 °C (left, top) or intravenously administered to a Sprague Dawley rat (left, bottom). Semi-logarithmic concentration vs time (right) demonstrating stability of XMc39 tracer miRNA in vitro (orange line) and in vivo (black line) over 240 min (n = 1).

**Figure 4:**
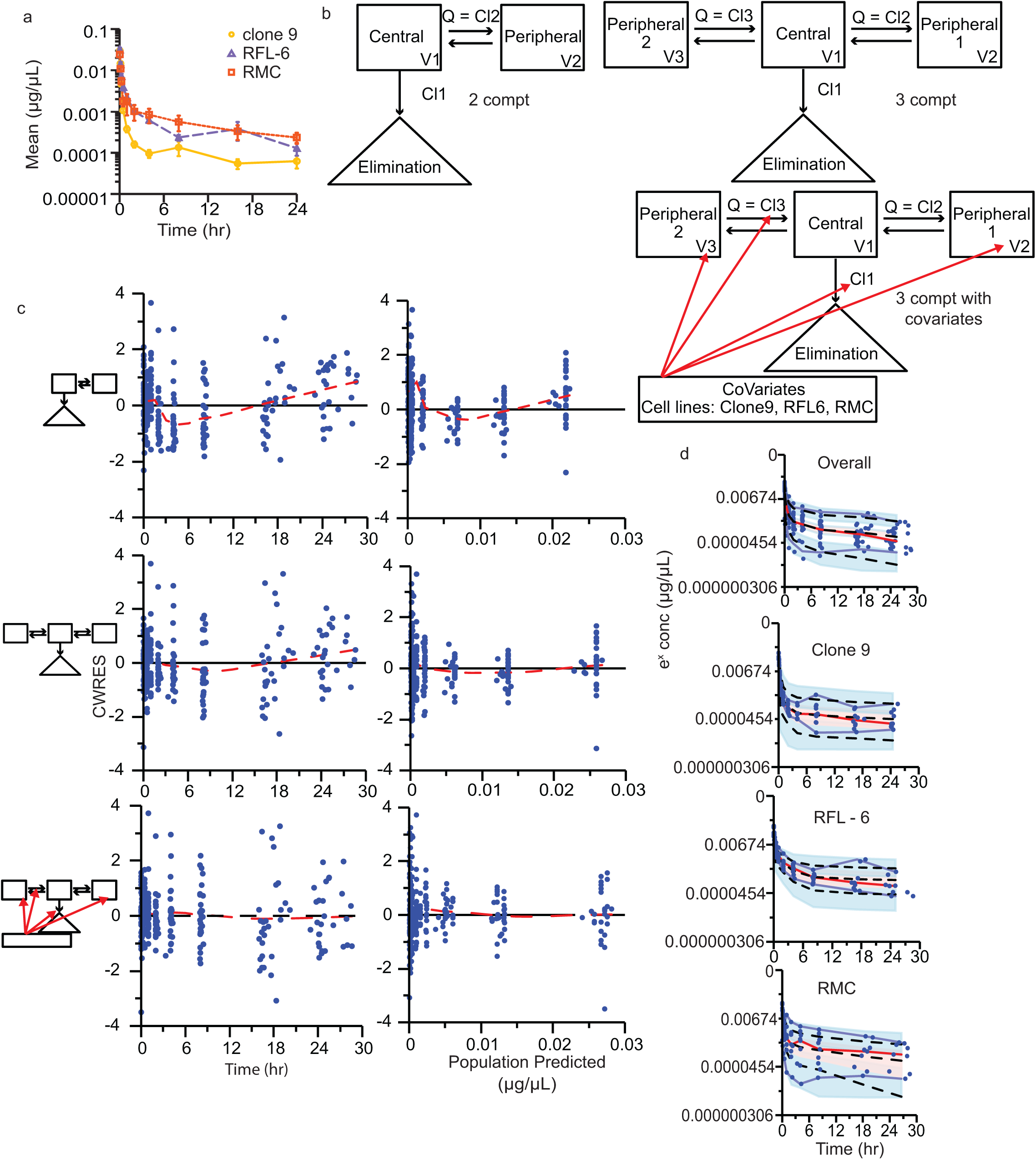
EV kinetic modeling. Final kinetic models for EVs administered to conscious Sprague Dawley rats. Each EV preparation had 9 - 10 animals (clone 9, n = 10; RFL-6, n = 9; RMC, n = 9). **a**, Mean normalized EV concentrations (± SE) over time after single intravenous bolus dose. Semi-logarithmic plot illustrating in vivo time course data for EVs isolated from clone 9 (yellow circles), RFL-6 (purple triangles), and RMC (orange squares) cell lines. **b**, Schematic representations of models: two-compartment (2 compt), three-compartment (3 compt), and three-compartment with covariates applied (3 compt with covariate). V = volume, Q = equal flow between two compartments (Phoenix software designates Q as numbered Cl parameters, e.g. Cl2 and Cl3). Red arrows indicate parameters to which the covariate was applied. **c**, Goodness-of-fit plots for conditional population weighted residuals (CWRES) vs. time and CWRES vs. population predicted concentrations. Model schematics are placed to the left of the respective plots. Blue circles indicate CWRES, black line indicates the zero residual reference line, dashed red line indicated the LOESS regression. See Supplementary Fig. 6 for additional Goodness-of-fit plots. **d**, Observation-based simulated posterior predictive evaluation with prediction-corrected visual predictive check (pcVPC) as semi-exponential concentration vs. time. From top to bottom, the plots include data from all cell lines, Clone 9, RFL-6, and RMC. Observed measures include individual observations (blue circles), median (dashed red line), and 5th and 95th percentiles (blue lines). Predicted measures include median, 5th, and 95th percentiles (dashed black lines). Shaded ribbons indicate predicted 90% confidence intervals around each quantile.

### Droplet digital PCR assay development and optimization

Ideally, pharmacokinetic studies are designed to accommodate five half-lives of the administered compound in order to measure approximately 97% of its elimination. We therefore required an assay with a dynamic range of five half-lives that was sensitive enough to detect very low-abundance tracer miRNA during terminal phase kinetics. Droplet digital PCR (ddPCR) is more sensitive than quantitative PCR^35^, and is capable of absolute copy number quantification instead of relative quantification against intra-assay standard curves. TaqMan stem-loop assays are preferred for the sensitive and specific detection of low-abundance miRNAs, but the additional 3’ exosome localization sequence in our XMc39 tracer rendered conventional cel-miR-39-3p assays unusable. Custom TaqMan assays require 3’ sequence specificity within 1-2 base pairs. We empirically determined the tracer miRNA sequence in labeled secreted EVs, and several variant 3’ sequences were identified (Fig. 2e). After consultation with the vendor, we designed a custom TaqMan assay, but we were unable to adequately detect tracer in EVs isolated from XMc39-transfected clone 9 cells (data not shown). We concluded that our tracer was incompatible with TaqMan chemistry. Hence, to quantify a heterogeneous population of tracer miRNA sequences, we designed an assay for use with the EvaGreen intercalating fluorophore. Conditions for ddPCR were optimized as follows.

The goal of ddPCR optimization is to maximize discrimination between positive and negative droplets (Fig. 3a) while also maximizing the sensitivity of target detection. Although EvaGreen is selectively fluorescent in the presence of double-stranded DNA molecules, it is sensitive to high concentrations of single-stranded DNA molecules such as primers. Using cDNA synthesized from a known quantity of purified RNA template, we optimized two critical parameters: primer concentration and annealing temperature (Ta) of the PCR reaction. Early optimization used a synthetic RNA oligonucleotide of the proprietary XMIR-NT sequence and corresponding forward primer, supplied by the vendor. Using a conservative starting primer concentration of 100 nM, we determined that 60 °C was the highest T_a_ to yield a discriminate band of positive droplets (Fig. 3b). Using 60 °C as an upper limit T_a_ to avoid non-specific amplifications that can occur at lower-than-optimal temperatures, we then tested a primer concentration gradient (Fig. 3c). The effect of primer concentration on EVAgreen fluorescence is evident in this figure, as higher primer concentrations clearly increase the median fluorescence of negative droplets. We determined the optimal primer concentration fell between 200-250 nM, based on band discrimination, percentage of positive droplets and tight clustering of positive and negative bands. Next, using a primer concentration of 250 nM, another T_a_ gradient was performed (Fig. 3d) which established 58 °C as optimal. A final primer concentration gradient (Fig. 3e) using this T_a_ indicated an optimal primer concentration of 200-225 nM. We determined 200 nM was preferable in order to minimize nonselective EvaGreen fluorescence. At this stage of development, the proprietary XMIR-NT sequence was replaced with cel-miR-39-3p (XMc39). Based on similarity in sequence length with the XMIR-NT forward primer, we used the same primer concentration of 200 nM to perform a final temperature gradient for XMc39 (Fig. 3f). We determined 56 °C was the optimal T_a_.

Commercial cDNA synthesis kits typically include single-stranded oligo(dT) adapters in the 2-5 μM range. These kits are not optimized for use with ddPCR, and the high amount of oligo(dT) carryover is enough to cause excessive EvaGreen fluorescence in droplets. We tested two strategies for minimizing oligo(dT) carryover into the ddPCR reaction. First, we varied the input amount of oligo(dT) by using an alternative “flex” cDNA synthesis kit from the same vendor which included separately packaged master mix components (data not shown). Second, we diluted the cDNA product 1:10 in ddPCR supermix and performed PCR preamplification immediately prior to droplet generation and ddPCR (Fig. 3g). The rationale for preamplification was twofold: lost sensitivity caused by the dilution of cDNA could be partially restored, and carryover oligo(dT) functions as a reverse primer that is consumed during preamplification. We found that both strategies minimized the “rain” (droplets that fall between negative and positive) (Fig. 3g). For applicability with the simpler non-“flex” protocol, we incorporated a 5 cycle preamplification into our standard protocol.

In order to establish linearity of the assay in a biologically relevant context, a synthetic RNA oligonucleotide representing the ideal XMc39 sequence was purchased and two-fold serial dilutions were prepared in miRNA extracted from naïve rat plasma. We then calculated expected ddPCR copy number values from known input amounts of oligonucleotide template and compared them to analytical data. As shown in Figure 3h (n = 3), the relationship between expected and observed copy numbers was highly linear (r^2^ = 0.997) and nearly identical (compared to the line of identity).

We noted during the course of these experiments that negative controls consisting of naïve plasma produced low, variable numbers of false positive droplets. To explore this, we prepared miRNA from two naïve plasma samples and analyzed replicate aliquots of each. These samples produced a random signal ranging from 20 to 266 copies with a CV of 110% (Fig. 3i). Negative controls consisting of water as the template for ddPCR amplification, however, consistently yielded no more than 1 positive droplet. Similar to the standard curve in Figure 3h, we analyzed lower concentrations of XMc39 and found that very low amounts of positive control RNA template reduced the number and variability of false positive droplets found in our negative controls (Supplementary Fig. 4). Taking this into consideration, we decided to use negative controls as a measure of quality control rather than a hard threshold for data exclusion. Sample sets with negative controls greater than 200 were reanalyzed using RNA as the starting material. Otherwise, we did not define a lower limit of quantification and allowed the model to use all the data.

### Retention and stability of EVs *in vivo*

To ensure that experimentally observed concentrations of tracer were not artifactual, we assessed for possible catheter contamination after dosing, as well as RNAse digestion of tracer in the blood. When dosing animals with highly concentrated EVs through catheters, laminar flow might cause EVs to be retained within the lumen. The interior volume of the jugular vein catheters used in this study was determined to be approximately 40 μL. We filled a catheter with 2 μg/μL labeled clone 9 EVs and then injected PBS in sequential 40 μL volumes, collecting the outflow each time. Less than 0.1% tracer was detectable in the 4^th^ sample relative to the 1^st^ (data not shown). We adopted the practice of using a pulsatile action when depressing the plunger of a syringe in order to create turbulent flow when flushing. Furthermore, we decided to use a volume of 500 μL when flushing the catheter with saline after dosing and 250 μL after sample collection.

Next, we performed a final validation of our EV preparation by testing the stability of incorporated tracer miRNA. Since blood plasma is rich in RNases that degrade unprotected circulating RNAs (data not shown), we needed to ensure that any observed elimination kinetics of tracer miRNA was specific to the behavior of its encapsulating vesicles and not due to RNAse degradation of free-floating molecules. For the stability assay, labeled EVs were intravenously administered to a live rat; in parallel, a proportional amount of the same labeled EV preparation was spiked into anticoagulated whole blood. The whole blood was incubated *in vitro* in a DNA LoBind tube at 37 °C. During the course of the experiment, *in vitro* blood samples were drawn immediately after *in vivo* blood samples at pre-specified time intervals (Fig. 3j). Tracer miRNA was stable for at least 4 h *in vitro*, whereas it exhibited marked elimination over the same 4 h *in vivo*. From this, we concluded that the detectable tracer miRNA in our EV preparations was protected from RNAse degradation in the blood and that observed *in vivo* kinetics are representative of the labeled EVs.

### *In vivo* kinetics of intravenously administered extracellular vesicles

This optimized method was applied to test our hypothesis that EVs from cultured cell lines of different origin exhibit different kinetics. Three Sprague Dawley-derived cell lines were selected for this study: clone 9 hepatocytes, RFL-6 lung fibroblast, and RMC mesangial cells. Liver, lungs, and kidneys have been identified as major organs of exosome clearance^36-40^. Labeled EVs were isolated from each of these cultured cell lines and intravenously administered to Sprague Dawley rats at a target bolus dose of 1,000 μg protein equivalent (range: 935-1,000 μg). EV preparations from each cell line were administered to 10 animals; thus, 30 animals were used in total. Blood samples were collected from each rat (clone 9, n = 10; RFL-6, n = 9; RMC, n = 9) at 2, 7.5, 15, 30, 60, 120, 240, 480, 960, and 1440 min following EV administration. Samples were processed in sets, analyzed by ddPCR, normalized to high and low standards between assays, and normalized to dose aliquots within assays (Supplementary Fig. 5, Supplementary Data 2). Two animals were excluded from analysis. One animal from the RFL-6 group was removed for concern of cross-sample contamination, and one animal from the RMC group for repeatedly failing quality control according to the pre-defined negative control threshold.

Normalized observed concentrations were plotted against the ideal collection time; differences between cell lines were visually apparent on a semi-log plot (Fig. 4a) and appeared to be multi-exponential, likely tri-exponential. Using ideal instead of actual collection times allowed for an analysis of standard error around the mean. Using industry standard pharmacokinetic modeling software, for compartmental analysis, we used first order conditional estimation - extended least squares (FOCE ELS) to estimate pharmacokinetic parameters. A one-compartment model would not execute in the modeling software. As reported in Table 1, a three-compartment model with one elimination from the central compartment (“3 compt model”) results in a much lower Akaike information criterion (AIC) value, which is a measure of model goodness of fit, than a two-compartment model with one elimination from the central compartment (“2 compt model”) (Table 1). Models with elimination from the central compartment are the simplest models^41^ (Fig. 4b), and likely exhibit the lowest AIC values because we only analyzed tracer miRNA concentrations in blood sampled from the central circulation.

**Table 1:** Model comparisons.

| Model | Akaike Information Criterion (AIC) |
| --- | --- |
| 1 compartment and 1 to 3 eliminations | Unable to estimate parameters |
| 2 compartment 1 elimination (central compt) | -3268.419 |
| 2 compartment 2 elimination | -3271.057 |
| 2 compartment 1 elimination (peripheral compt) | -1864.035 |
| 2 compartment 3 elimination (2 central, 1 peripheral compt) | -3260.423 |
| 3 compartment 1 elimination (central) | -3324.393 |
| 3 compartment 1 elimination (central) - cell line covariate applied to all parameters | -3364.679 |
| 3 compartment 1 elimination (central) – cell line applied to V2, V3, Cl, Cl3 | **-3367.219** |
| 3 compartment with 2 elimination (both peripheral) | -3313.1419 |
| 3 compartment with unequal distribution rates between compt and 1 elimination (central) | -3328.015 <sup>a</sup> |
| 3 compartment – two unequal distribution rates between compt - 1 elimination (central)<br>Cell line covariate applied to all parameters | -3321.4789 <sup>a</sup> |
| 3 compartment 3 elimination | -3312.0896 |

a – No precision estimates for parameters

Covariates of cell line, weight, and batch were incorporated into the three-compartment model, and only the cell line covariate resulted in a meaningful decrease in the AIC value and change in eta-covariate comparisons. Our protocol required animals to fall within a narrow weight range, so it is not surprising that weight had little effect as a covariate. Using a shotgun approach of applying the cell line covariate to each parameter, we found that applying the covariate to Volume 2 (V2), Volume 3 (V3), Clearance (Cl), and Clearance 3 (Cl3) (Fig. 4b) resulted in the lowest AIC value (Table 1). Code for execution of the model can be found in Supplementary Note.

### Population model evaluation

We compared goodness of fit scatterplots between the two- and three-compartment models (Fig. 4c; Supplementary Fig. 6). Goodness of fit plots evaluate the population residual error created from population predictions subtracted from the observations. Ideally, the weighted residual error should have a mean of zero (horizontal line) and randomly distribute around the horizontal line (variance). Model fitness is improved as the LOESS regression line approaches the ideal weighted residual line of zero. In evaluating the conditional population weighted residual (CWRES) versus time and versus the population predicted concentrations, the three-compartment model improved the model fit of the data (Fig. 4c). In evaluating observed concentrations versus individual predicted concentrations and population predicted concentrations, the LOESS regression line approached the line of unity indicating that the three-compartment model again outperformed the two-compartment model (Supplementary Fig. 6). The addition of the covariates to the three-compartment model further improved upon the base model (Fig. 4 c, Supplementary Fig. 6) with individual model fits in Supplementary Fig. 7.

We performed an observation-based simulated posterior predictive evaluation with prediction-corrected visual predictive check (pcVPC, Fig. 4d). The pcVPC used a log-additive error model to prevent simulating negative concentrations. The pcVPC simulates concentration data from the three-compartment model with covariates and plots the distribution of observations and the distribution of the predicted concentrations over time. The simulated three-compartment model with covariates contains the observed data within the shaded confidence interval, which suggests a good model description.

### Model outcome and performance

Notably, the volume of distribution for the central compartment (28 mL) is similar to the mean calculated total blood volume of a male Sprague Dawley rat^18^ with an average weight of 372 ± 6 g, or 26 ± 0.4 mL (mean ± S.E.). As shown in Table 2, the half-life of elimination ranged from 12 h to 215 h across the 3 cell lines and was significantly different between them. The volume of distribution between the central compartment and first peripheral compartment was significantly different between the clone 9 and RFL-6 cell lines. The volume of distribution between the central compartment and second peripheral compartment was significantly different between all cell lines. The area under the concentration-time curve (AUC) was significantly different between RMC and clone 9, and between RMC and RFL-6.

**Table 2:**
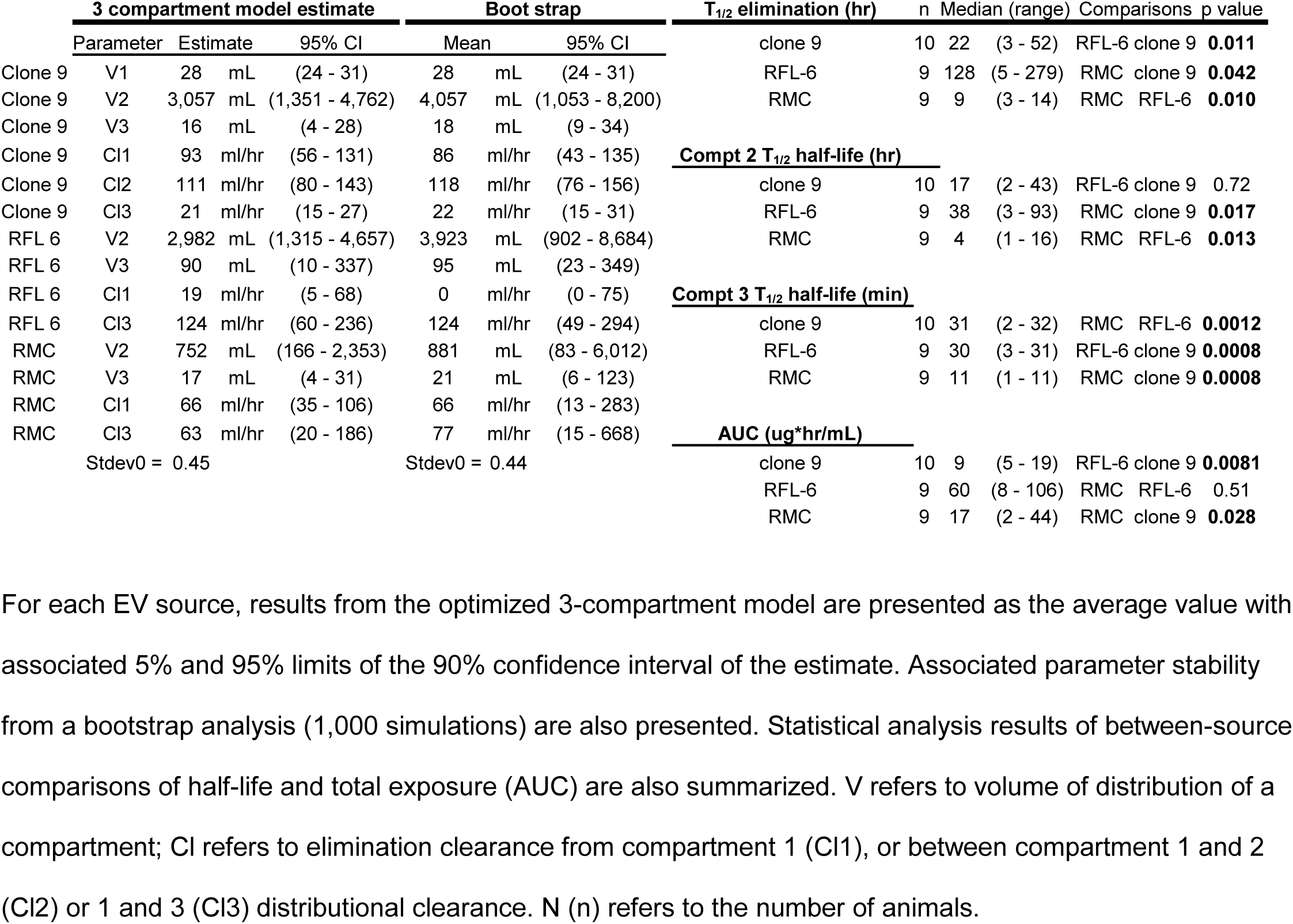
Population pharmacokinetic parameter estimates describing plasma exosome exposure following intravenous administration.

A bootstrap analysis using 1,000 simulations was performed to evaluate the likelihood of achieving similar results if the experiment was replicated. Overall, the bootstrapped results mirrored the actual experiments. One exception is that the clearance of elimination from the central compartment (Cl) was similar for all 3 cell lines (Table 2, Supplementary Fig. 8). This suggests that cell line differences in EV kinetics are due to differences in EV distribution to the peripheral compartments.

## Discussion

EVs continue to attract broad interest as both targeted therapeutics and dynamic biomarkers in the systemic circulation, yet modalities for the study of *in vivo* EV kinetics are limited to modifications of membrane composition that provide a partial picture of how composition affects kinetics. Here, we detail an accessible method for measuring the *in vivo* kinetics of EVs derived from cultured cells. Conceptually, we integrate several major techniques in this approach. First, an expression vector is used to encode a non-homologous tracer miRNA that is selectively packaged into exosomes. Next, labeled EVs are harvested from enriched cell culture media and injected into rats. Lastly, ddPCR is used for the sensitive detection of cDNA prepared from low-volume blood samples. Compared to other reported techniques^4-6^, our method bypasses the need for modifying EV surfaces with external ligands. We also designed our protocol so that an entire kinetic time course may be performed using a single animal, minimizing the variability of pooling different time points across multiple animals.

We successfully applied our method to test a hypothesis that EVs from different cell lines exhibit different kinetics *in vivo*. These studies quantitatively described for the first time, though with important caveats, significant differences in kinetic parameters between EVs derived from liver, lung, and kidney cells. The known biology of EVs suggests that multiple routes of elimination are likely to exist (e.g. tissue sequestration, intracellular degradation, and excretion). We did not have enough data to support a model with elimination from a peripheral compartment, however a three-compartment model supports the idea that EVs circulate in the vasculature and then move between shallow and deep peripheral compartments potentially representing tissue distribution and intracellular metabolism.

A three-compartment model best described the observed kinetics of EVs derived from all three cell lines used in this study. While there was a high degree of reproducibility between EV preparations from multiple passages of the same cell line, one caveat is that differences between cell lines should be interpreted with some caution. Cell lines were grown under optimal conditions recommended by ATCC, which necessitated the use of different base culture mediums and different concentrations of FBS. While it is possible that differences in nutrient and serum concentrations might generally influence EV composition, and thus kinetics, our intention was to preserve the intended characteristics of each cell line without subjecting them to the possible stress of suboptimal culture conditions.

As for the second caveat, we must note that the polyethylene glycol (PEG)-based EV isolation method used in our study, while often chosen by researchers for its relative ease of use compared to ultracentrifugation-based methods, is known to provide high yield at the expense of purity^25,29,42^. Using standard protocols, PEG isolation is best described as providing a crude source of sEVs with the possibility of co-isolating non-EV contaminants. Since this was the first application of our method to measure *in vivo* kinetics of EVs, we chose to use PEG isolation of EVs for its two main advantages: high yield and reproducibility across studies. In order to minimize the influence of potential contaminants, three steps were included in addition to the standard protocol. First, we subjected EV-enriched media to an additional centrifugation step in order to remove larger microvesicles and small debris^23,24^. Second, the EV pellet was gently washed to remove residual PEG prior to resuspension. Third, the resuspended EV pellet was passed through a size exclusion column to remove small molecules such as unbound oligonucleotides and residual PEG. The resulting size distributions, transmission electron micrographs, and western blots are representative of sEV preparations using other methods^37,38,40,43^, and lipidomic characterization of our EV preparations provides further support by demonstrating enrichment of endosomal lipids. Interestingly, the EVs prepared for this study did not induce any noticeable inflammatory response in recipient animals; this may be due to the utilized cell lines originating historically from Sprague Dawley rats.

The expression vector used in our study appends a localization motif to the encoded stem-loop tracer miRNA sequence, and is expected to selectively enrich EVs of endosomal origin with the mature sequence^19^. Although we demonstrated that tracer miRNA in our EV preparations was resistant to RNAse degradation and took this as evidence for its encapsulation within vesicles, our chosen method of EV isolation implicates two miRNA-binding proteins, Ago2 and hnRNPA2/B1, as possible co-isolated contaminants. Ago2 protects bound miRNA from RNAse degradation, evidenced by the stability of non-vesicular circulating miRNAs in plasma^44^, but evidence for hnRNP proteins conferring similar resistance is less convincing^45^. Ago2-bound miRNA is abundant in plasma^46^ and has been detected with EVs isolated by PEG from plasma^27^, but several studies have failed to identify Ago2 in conditioned media^47^ or EVs prepared from cultured cell lines^47-50^, with two exceptions^33,51^. These differences in observations may be attributable to any number of experimental conditions, including the type of cell lines used. We assayed our EV preparations by immunoblotting with a pan-Argonaute (Ago1-4) antibody and found variable amounts of protein, including distinct bands in two of three of our EV samples (Fig. 2). In the case that co-precipitated protein-bound miRNA may have resulted in a measurable signal, we have no reason to believe that the proteins of greatest concern (e.g. Ago2, hnRNPs) would cause cell-specific differences in elimination kinetics, nor introduce significant artifacts in our model that falsely attribute differences in elimination kinetics to the cell line covariate. However, since the method presented in this report is applicable to measuring the kinetics of EVs isolated by any method, including ultracentrifugation, it would indeed be possible to quantify the influence of different EV isolation methods on EV kinetics *in vivo* and to determine the best-suited methods for such characterizations.

In this report, we demonstrate how the rich, quantitative data obtained by our method may be used with nonlinear mixed-effects modeling to produce kinetic models of EVs secreted from cultured cells *in vitro* and administered *in vivo*. We have detailed the necessary points to consider when developing such an assay, including good practices for establishing ddPCR assay conditions and analytical reproducibility. One limitation of our study is the inability of our biological negative control (naïve plasma) to accurately define a lower limit of quantification. As demonstrated, there was higher variability of false positive droplets in samples prepared from naïve plasma than from samples prepared using very dilute positive controls. This suggests random off-target amplification that is reduced when target template is present, even at very low concentrations.

Our findings demonstrate the ability of our method to parse the kinetic differences between EVs isolated from different cellular origins or grown under different conditions. We expect this to be a powerful tool for developing systems pharmacology-based and physiologically-based EV kinetic models that elucidate heretofore unknown characteristics of EV biology. This will make it possible to develop and use *in vivo* kinetic models to discover EV function and better design studies for therapeutics and circulating biomarkers.

Aspects of our method may be easily modified to explore different biological questions. For instance, EVs of cultured cells may be interrogated for differences in response to various treatments, compared to untreated controls. Additionally, labeling and vesicle isolation protocols may be altered to study other EVs such as microvesicles and apoptotic bodies. Ultracentrifugation-based methods may be used to further isolate subpopulations of EVs. By using conventional techniques and reagents, our method can be tailored to address a variety of scientific questions.

## Funding Statement

This research was supported by the Department of Medicine, Indiana University School of Medicine, Indianapolis, IN; and the National Institutes of Health-National Institute of General Medical Sciences under Grant K08GM119006.

### Acknowledgements

T.D. thanks the following people: Dr. Benson, for financial assistance and mentorship in this investigation; Dr. R. Graham Cooks at Purdue University for oversight of lipidomic mass spectrometry; Katherine A. Hargreaves for assistance with animal surgeries; Kari McClimon for training in jugular vein catheterizations.

## Author Contributions

E.A.B. conceived the study. E.A.B. and T.D. designed the experiments. T.D. performed the animal surgeries with assistance from E.A.B. T.D. performed all experiments and assay development with E.A.B.’s supervision. E.A.B. curated the data and performed pharmacokinetic analysis and compartmental modeling with supervision and guidance by R.E.S. M.E.E. performed MS data acquisition, assisted in developing the MRM profiling method, and authored mass spectrometry methods under C.R.F.’s supervision. T.D. analyzed MS data with instruction by C.R.F. E.A.B. and T.D. wrote and edited the manuscript with input from the other authors. All authors approved the final manuscript.

