## Supplementary Figures for "Quantification of extracellular vesicles with unaltered surface membranes using an internalized oligonucleotide tracer and droplet digital PCR applied to modeling of *in vivo* kinetics"

Table of Contents:

Supplementary Fig. 1: XMIRXpress plasmid map
Supplementary Fig. 2: EV size distribution based on cell line of origin
Supplementary Fig. 3: Principal component analysis of 31 top differentiating EV lipids between cell lines
Supplementary Fig. 4: Negative control plasma does not represent the lower end of quantification
Supplementary Fig. 5: Diagram and schematic of conversion factors used to normalize ddPCR copy numbers to EV concentration
Supplementary Fig. 6: Goodness-of-fit plots, observed concentration vs. individual and population predicted concentrations (IPRED, PRED)
Supplementary Fig. 7: 3 compartment with covariate EV plots for every subject's time course
Supplementary Fig. 8: Box plots of the overall half-life of elimination, overall AUC, and distributional half-lives between the central and peripheral compartments in the final 3 compartment with covariates model
Supplementary Fig. 9: Unedited western blot of CD63 (non-reduced conditions)
Supplementary Fig. 10: Unedited western blot of CD63 (reduced conditions)
Supplementary Fig. 11: Unedited western blot of CD81 (non-reduced conditions)
Supplementary Fig. 12: Unedited western blot strip of CD81 (reduced conditions)
Supplementary Fig. 13: Unedited western blot of tsg 101
Supplementary Fig. 14: Unedited western blot of Alix
Supplementary Fig. 15: Unedited western blot of ApoA-I
Supplementary Fig. 16: Unedited western blot of Histone H3.1
Supplementary Fig. 17: Unedited western blot of Cyt c
Supplementary Fig. 18: Unedited western blot of GM130
Supplementary Fig. 19: Unedited western blot of α-actinin
Supplementary Fig. 20: Unedited western blot of Ago1-4
Supplementary Fig. 21: Unedited western blot of hnRNP A2/B1

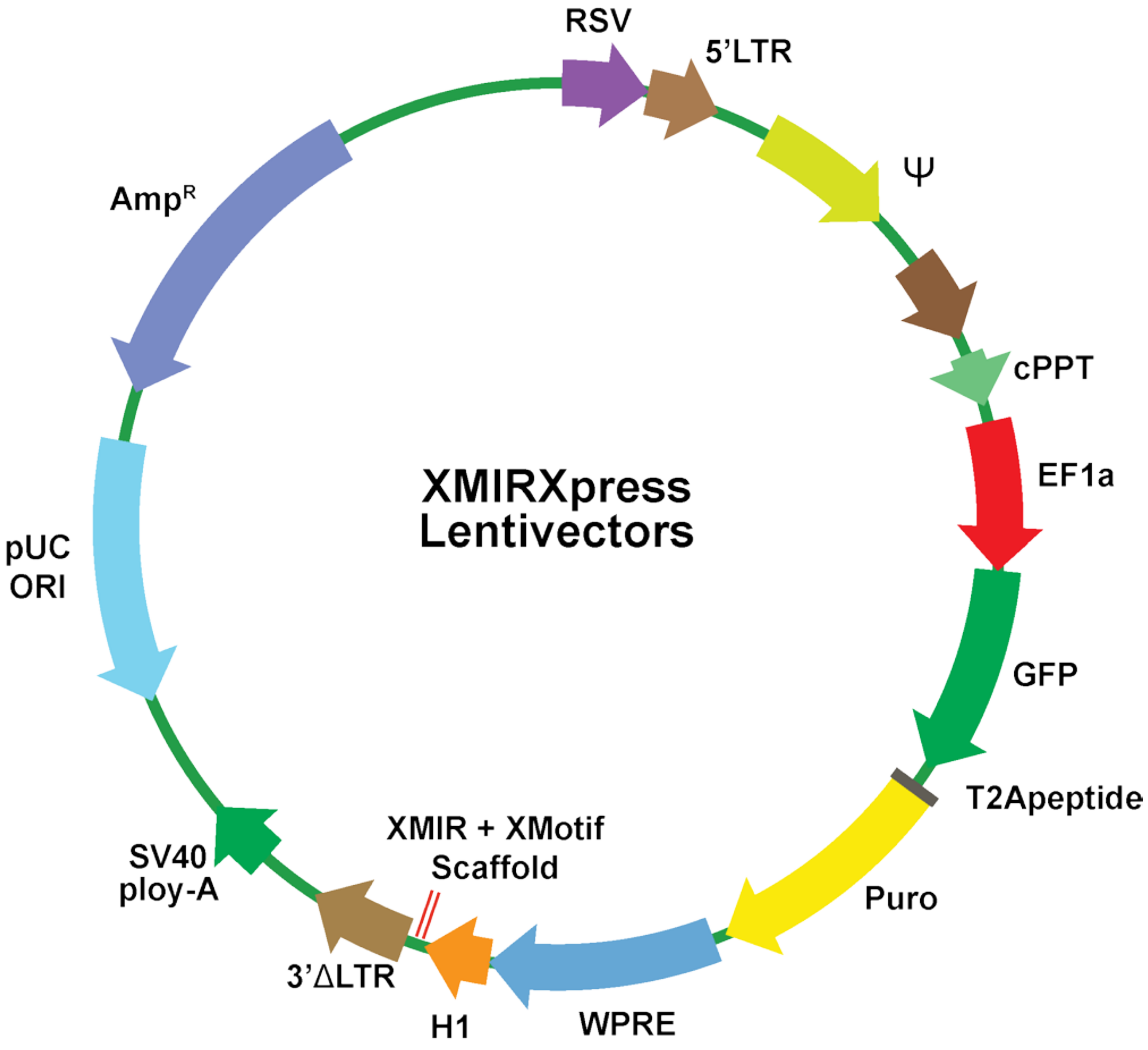


**Supplementary Fig. 1: XMIRXpress plasmid map**

XMIRXpress lentivector plasmid which contains the XMotif exosome localization signal. Double red lines indicate the site of tracer miRNA insertion. $AMP^R$ – ampicillin resistance, pUC ORI – origin of replication, RSV 5'LTR – rous sarcoma virus long terminal repeat promoter, ψ – sequence for viral packaging of RNA into capsid, cPPT – central polypurine tract to enhance transduction, EF1a-GFP-Puro – promoter for green fluorescent protein and puromycin resistance, WPRE – woodchuck hepatitis virus post-transcriptional regulatory element, H1 – polymerase III promoter, 3' delta LTR – for self-inactivation of lentivirus vector by terminating transcription initiated by 5' LTR, SV40 poly-A – secondary transcription termination.

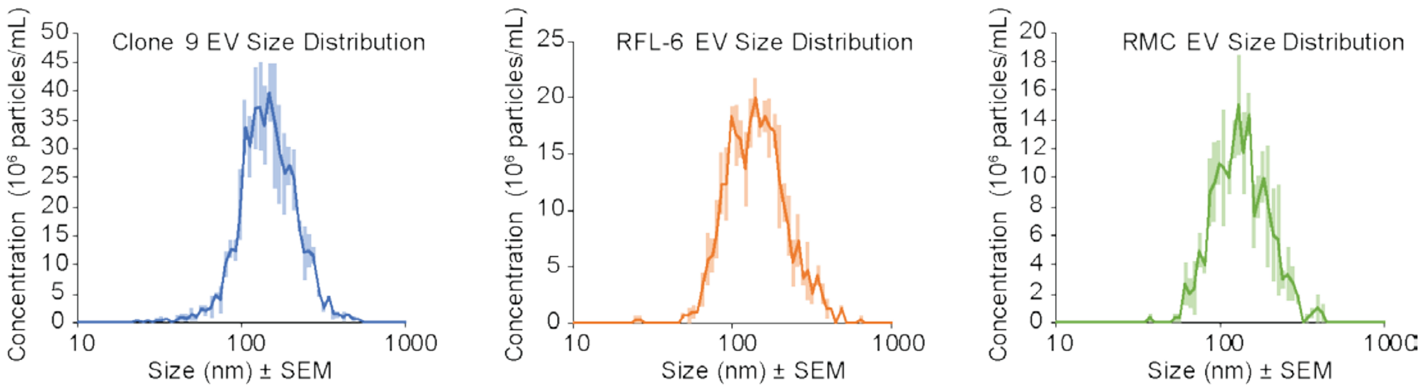


**Supplementary Fig. 2: EV size distribution based on cell line of origin**

Average size distributions of EVs from cultured clone 9 hepatocytes, RFL-6 lung fibroblasts, and RMC mesangial kidney cells (n = 3 for each cell line, mean ± S.E.). Bin counts are represented as absolute concentrations for each cell line as opposed to relative proportion.

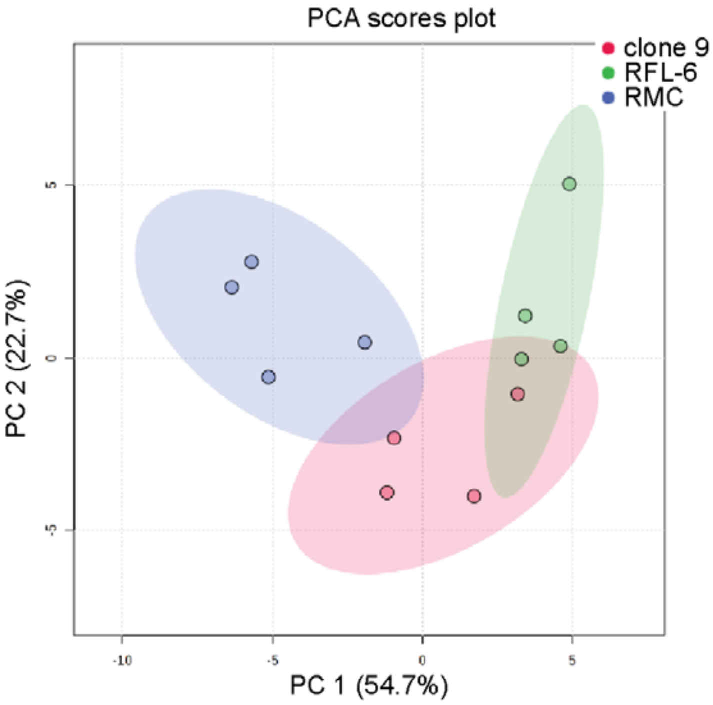


**Supplementary Fig. 3: Principal component analysis of 31 top differentiating EV lipids between cell lines.**

Scores plot for the selected principal components (PC1 and PC2). Explained variances are shown in parentheses on the axes. There were 4 replicates per cell line; clone 9 in red, RFL-6 in green, and RMC in blue.

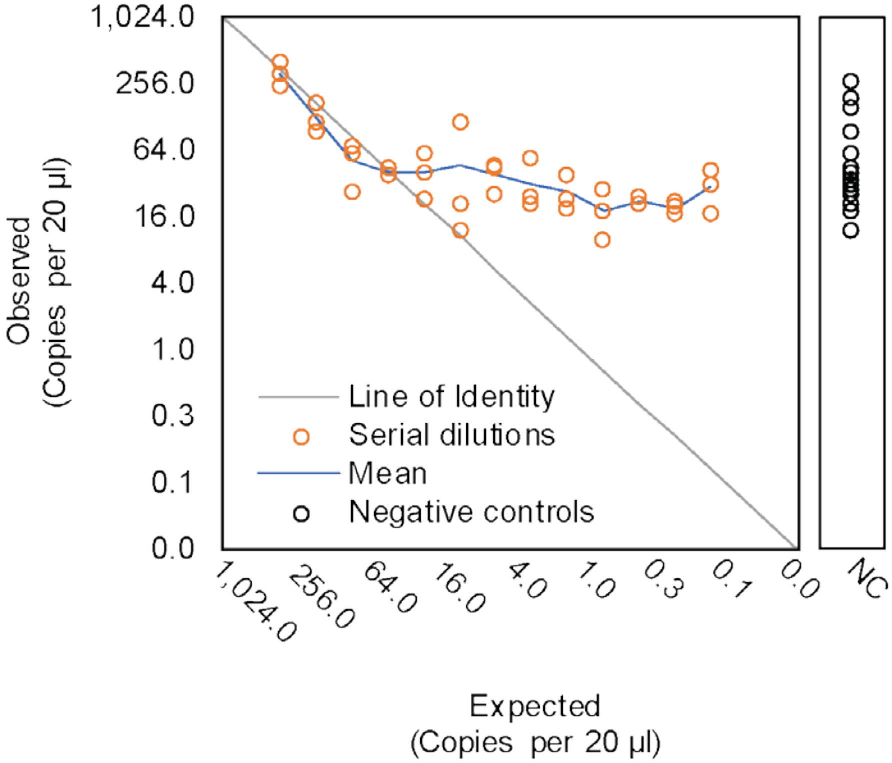


**Supplementary Fig. 4: Negative control plasma does not represent the lower end of quantification**

Two-fold serial dilutions of XMc39 RNA plotted as observed vs. expected copies/20 µL. Adjacent to the main graph is a plot of observed negative control copy concentrations (n = 15 technical replicates from 2 biological replicates, also plotted in Fig. 3i), using the same y-axis values as the main graph. Individual observed concentrations (orange circles), mean (blue line), line of identity (dashed gray line), and negative controls (NC, black circles).

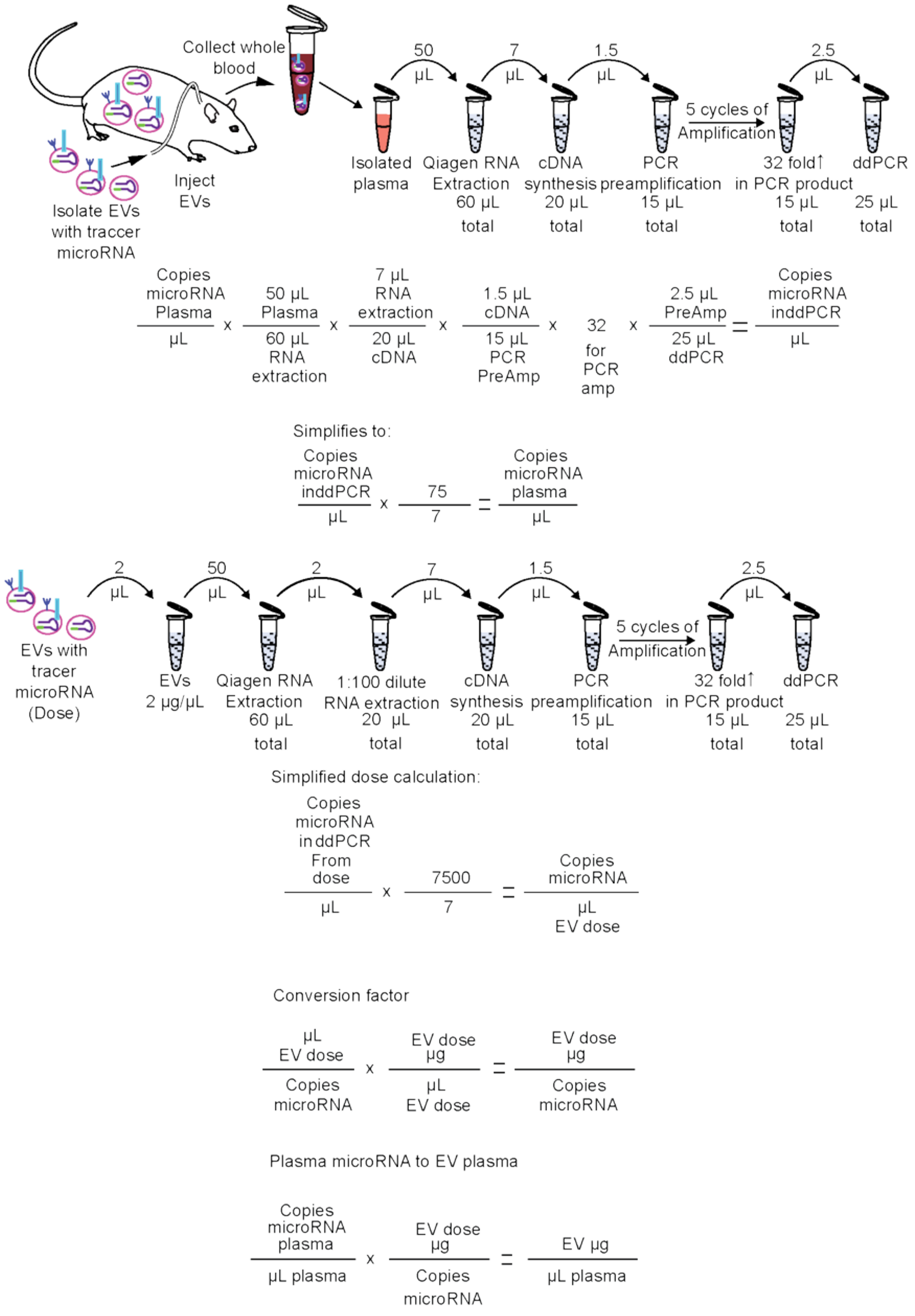


**Supplementary Fig. 5: Diagram and schematic of conversion factors used to normalize ddPCR copy numbers to EV concentration**

**Top schematic** illustrates all the steps and volumes necessary to obtain the final tracer (microRNA) concentrations in ddPCR samples. Accompanying equations constitute the conversion of microRNA concentration in ddPCR samples to the concentration of microRNA in originating plasma. **Bottom schematic** illustrates all the steps and volumes necessary to obtain microRNA concentrations in dose aliquots. Accompanying equations provide the conversion to microRNA concentration in ddPCR samples to microRNA concentration in the dose. Final equations provide calculations for the transformation of miRNA concentrations to EV concentrations (as units of protein equivalent).

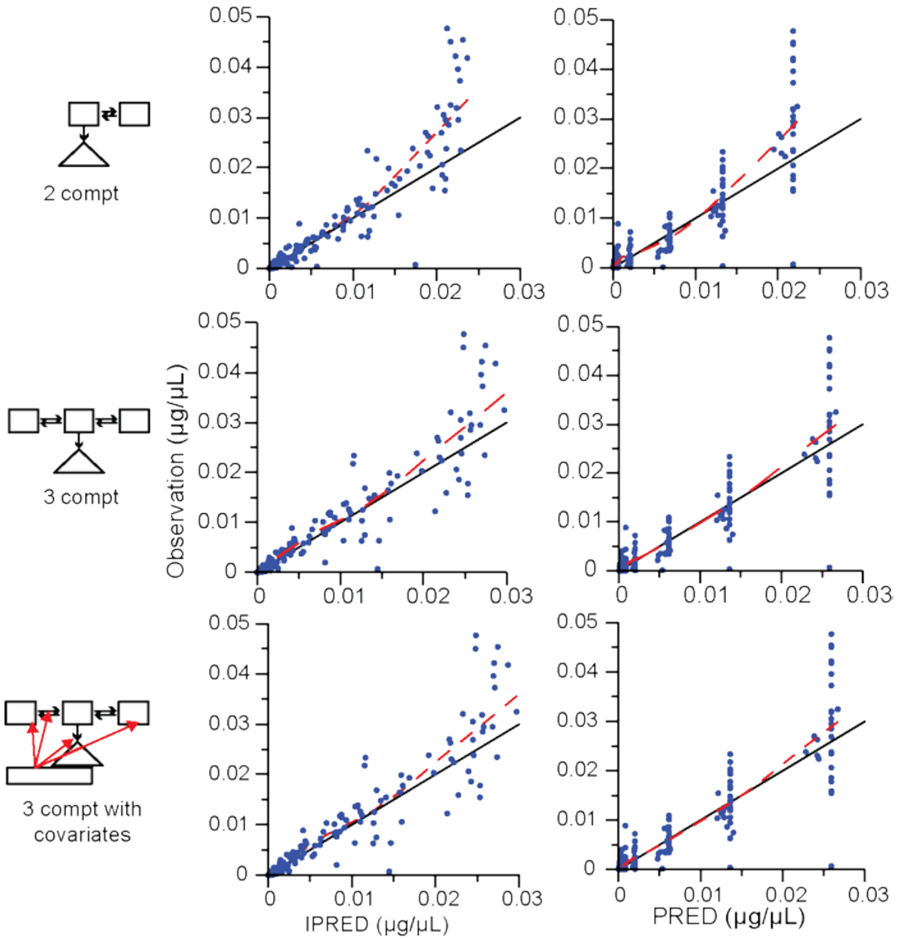


**Supplementary Fig. 6: Goodness-of-fit plots, observed concentration vs. individual and population predicted concentrations (IPRED, PRED)**

Blue dots are the observed vs. IPRED (left side graphs) or PRED (right side graphs) concentrations, black line is the line of identity, dashed red line indicates the LOESS regression for 2 compt, 3 compt and 3 compt with covariates models.

Observations, population predictions, individual predictions vs time

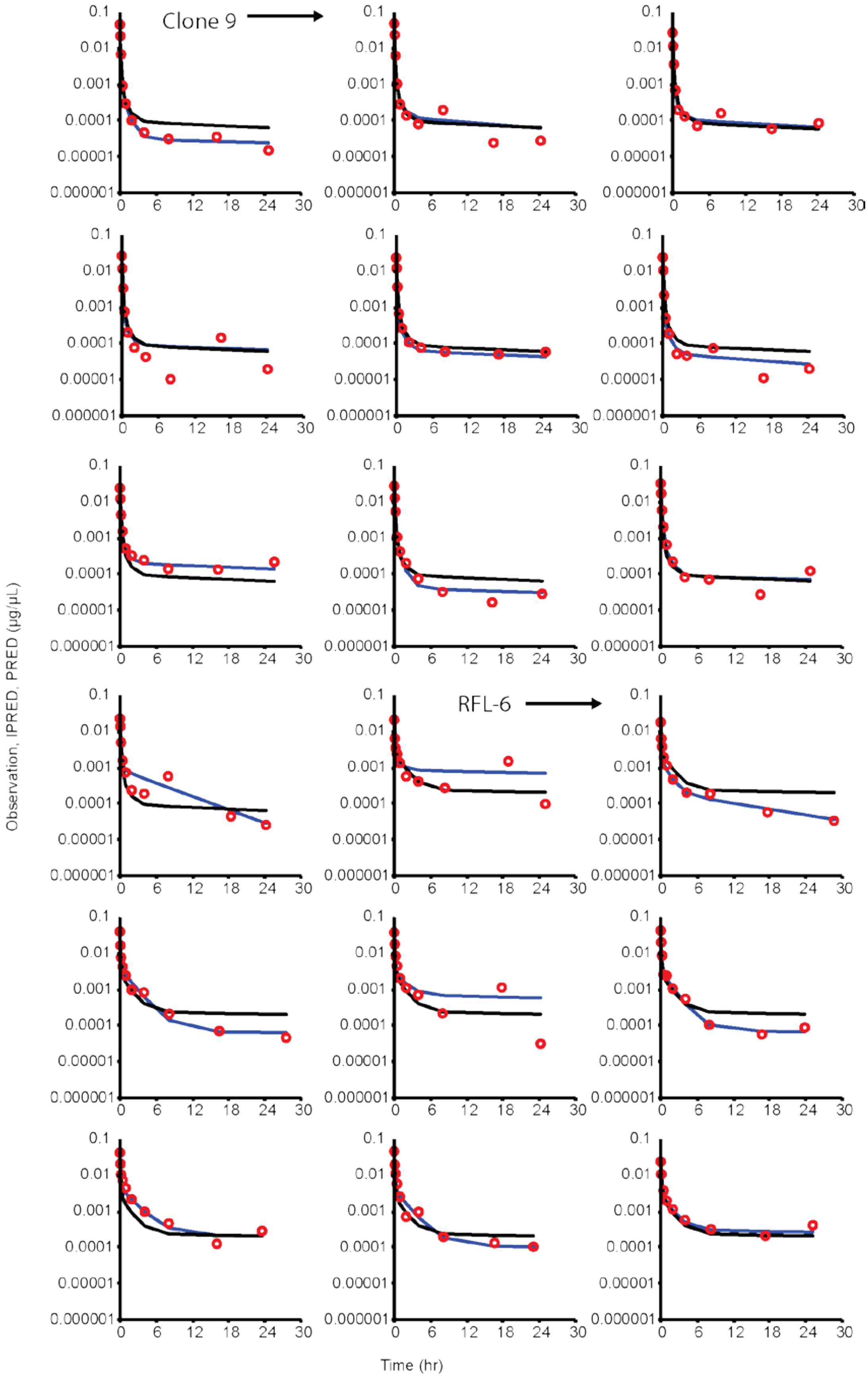


*continued…*

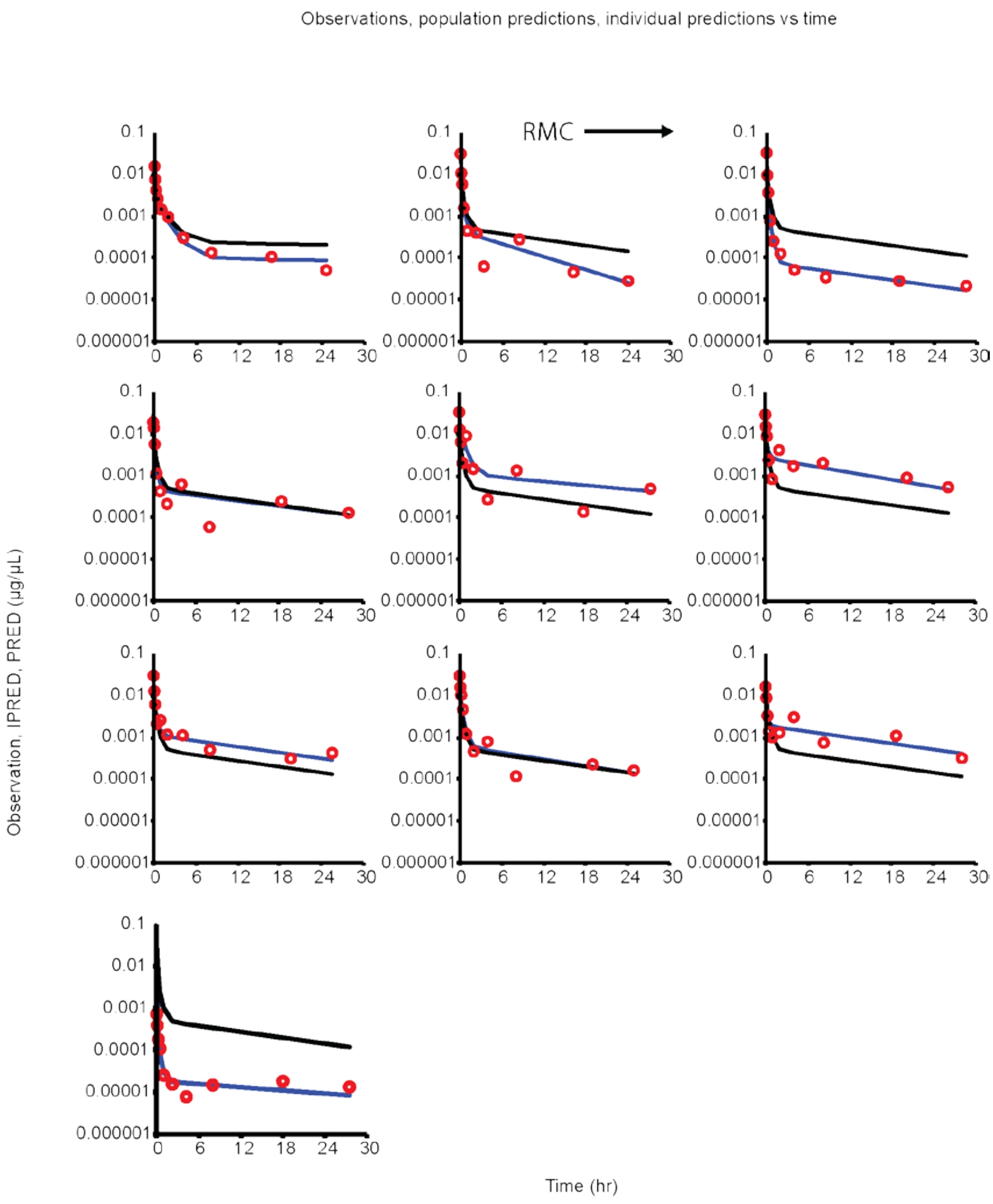


**Supplementary Fig. 7: 3 compartment with covariate EV plots for every subject's time course.**

Semi-logarithmic plots of the observed (red circles), population predicted (PRED, black lines) and individual predicted concentrations (blue lines) over time in each animal subject using the final 3 compt model with cell line covariate. In order, as indicated by arrows: clone 9 (n = 10), RFL-6 (n = 9), RMC (n = 9). All concentrations are in µg/µL.

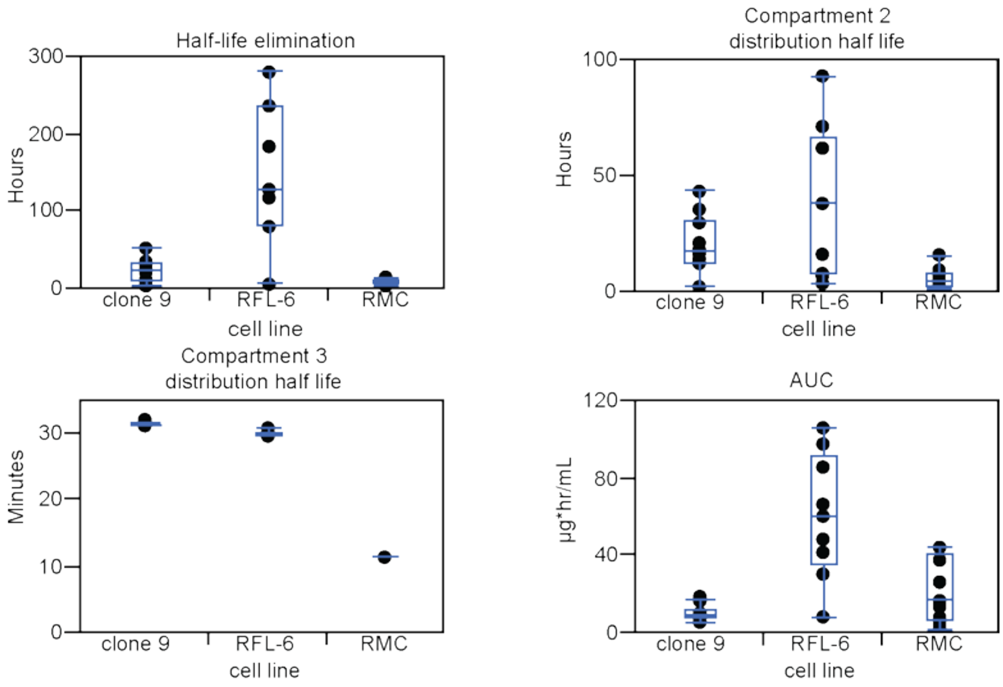


**Supplementary Fig. 8: Box plots of the overall half-life of elimination, overall AUC, and distributional half-lives between the central and peripheral compartments in the final 3 compartment with covariates model.**

Box plots for each cell line for the data presented in Table 2. Blue box indicates 25%, 50% (median), and 75% quartiles, whiskers are the maximum and minimum. There are no outliers in the data.

a
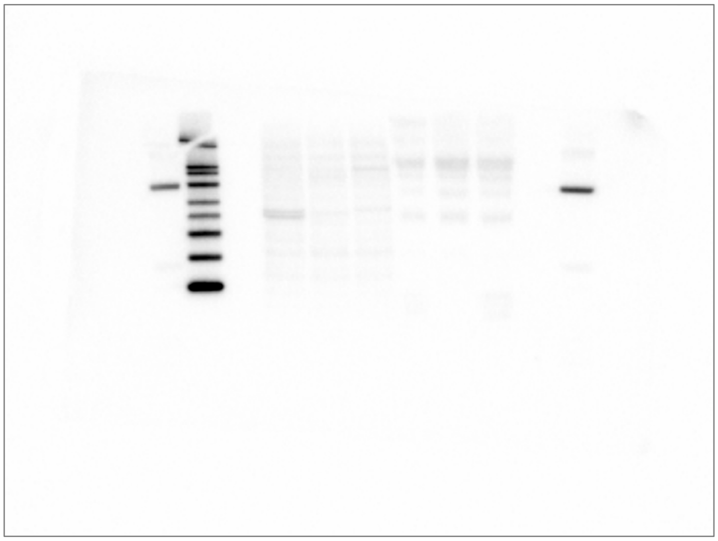

b
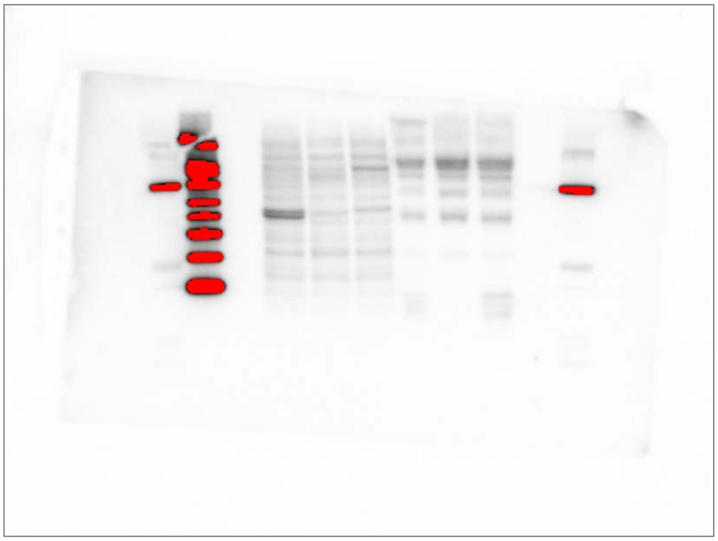

c
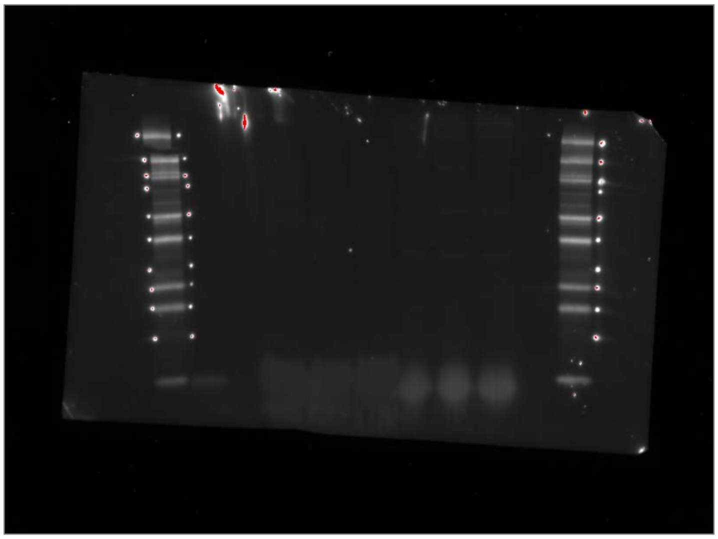

**Supplementary Fig. 9: Unedited western blot of CD63 (non-reduced conditions).**

Columns from left-to-right: Kaleidoscope Protein Standards, MagicMark Protein Standards, Empty Lane, 50 μg Clone 9 WCL, 50 μg RFL-6 WCL, 50 μg RMC WCL, 50 μg Clone 9 EV, 50 μg RFL-6 EV, 50 μg RMC EV, Empty Lane, Kaleidoscope Protein Standards.

For chemiluminescent detection of HRP, image acquisition was optimized for both (a) intense bands and (b) faint bands. Fluorescent detection (c) of the Kaleidoscope Protein Standards was performed using settings for Alexa Fluor 680 and optimized for intense bands. Red color represents signal saturation. Dots next to protein standards in the fluorescent image are a byproduct of labeling the visible standards with a laboratory marker.

a b

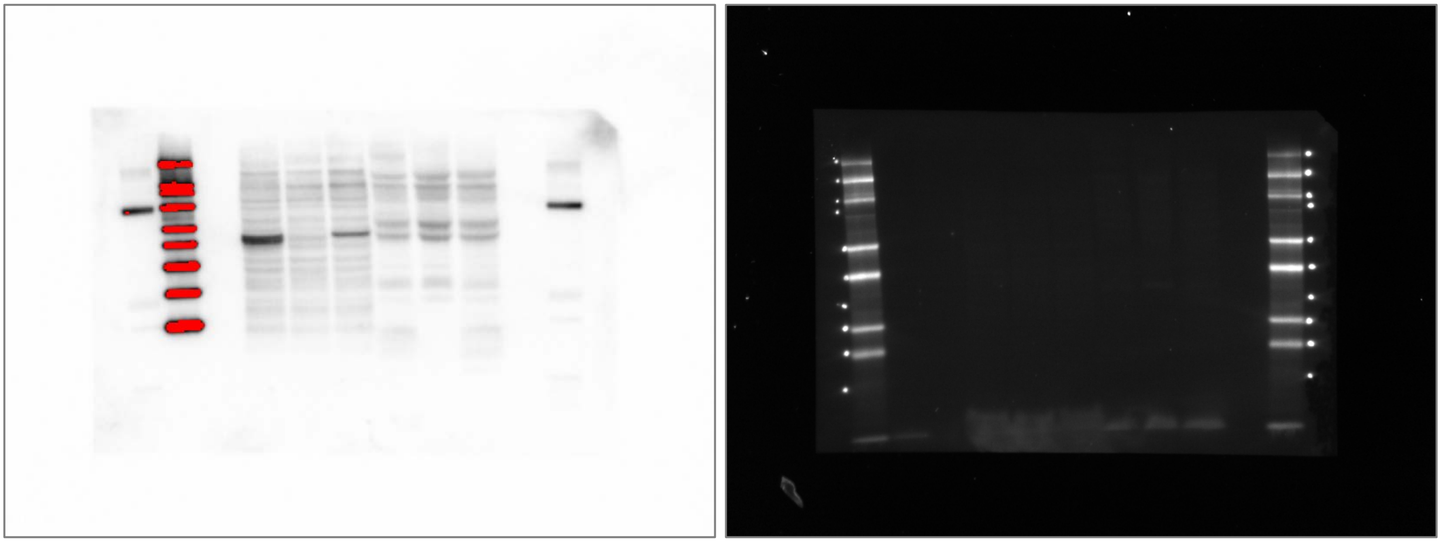

**Supplementary Fig. 10: Unedited western blot of CD63 (reduced conditions).**

Columns from left-to-right: Kaleidoscope Protein Standards, MagicMark Protein Standards, Empty Lane, 50 μg Clone 9 WCL, 50 μg RFL-6 WCL, 50 μg RMC WCL, 50 μg Clone 9 EV, 50 μg RFL-6 EV, 50 μg RMC EV, Empty Lane, Kaleidoscope Protein Standards.

For (a) chemiluminescent detection of HRP, image acquisition was performed using an exposure time of 1.5 seconds. Fluorescent detection (b) of the Kaleidoscope Protein Standards was performed using settings for Alexa Fluor 680 and optimized for intense bands. Red color represents signal saturation. Dots next to protein standards in the fluorescent image are a byproduct of labeling the visible standards with a laboratory marker.

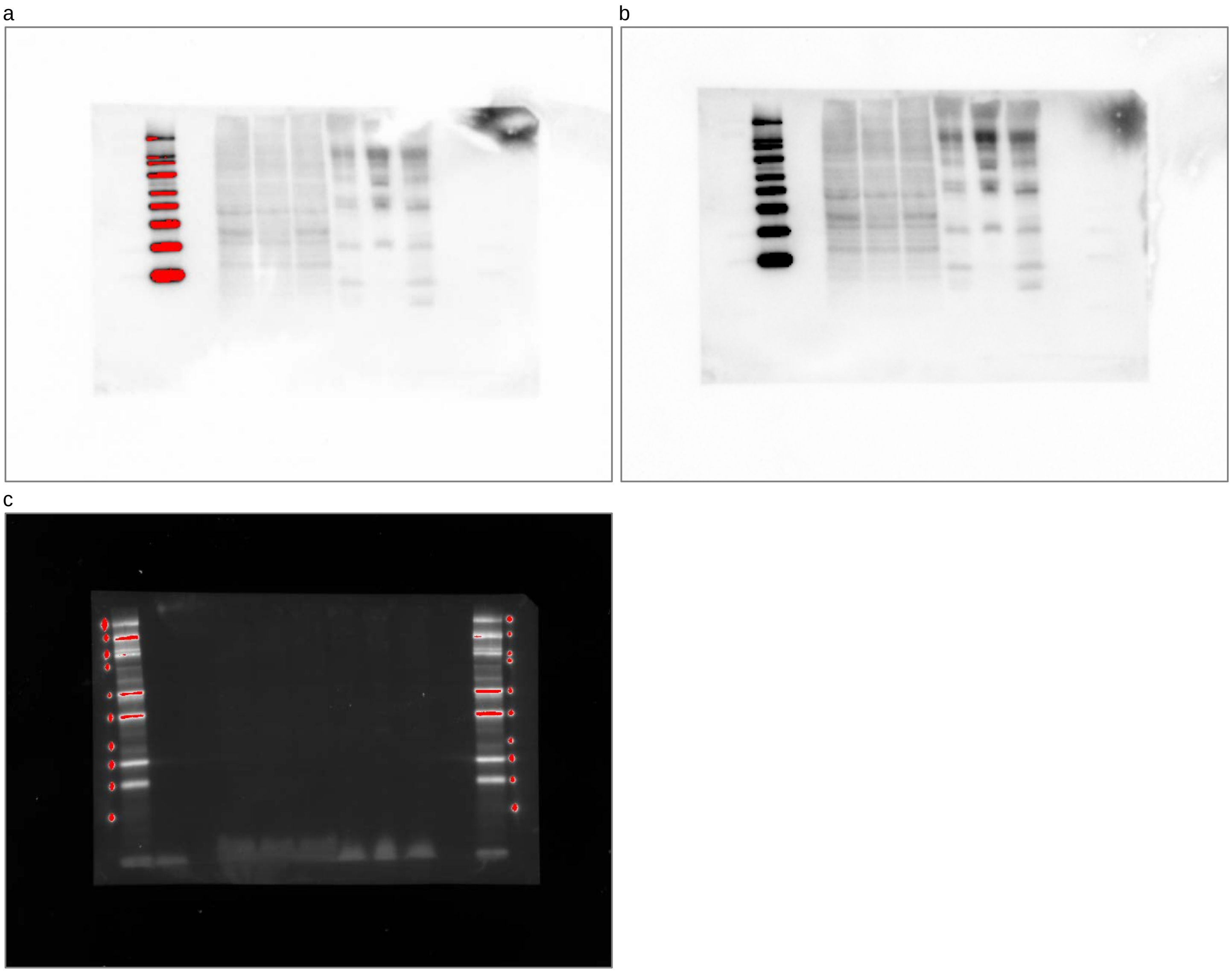


**Supplementary Fig. 11: Unedited western blot of CD81 (non-reduced conditions).**

Columns from left-to-right: Kaleidoscope Protein Standards, MagicMark Protein Standards, Empty Lane, 50 μg Clone 9 WCL, 50 μg RFL-6 WCL, 50 μg RMC WCL, 50 μg Clone 9 EV, 50 μg RFL-6 EV, 50 μg RMC EV, Empty Lane, Kaleidoscope Protein Standards.

For chemiluminescent detection of HRP, image acquisition was optimized for (a) faint bands, then (b) reacquired using a 10 second exposure time. Fluorescent detection (c) of the Kaleidoscope Protein Standards was performed using settings for Alexa Fluor 680 and optimized for intense bands. Red color represents signal saturation; this option was turned off when acquiring (b). Dots next to protein standards in the fluorescent image are a byproduct of labeling the visible standards with a laboratory marker.

a

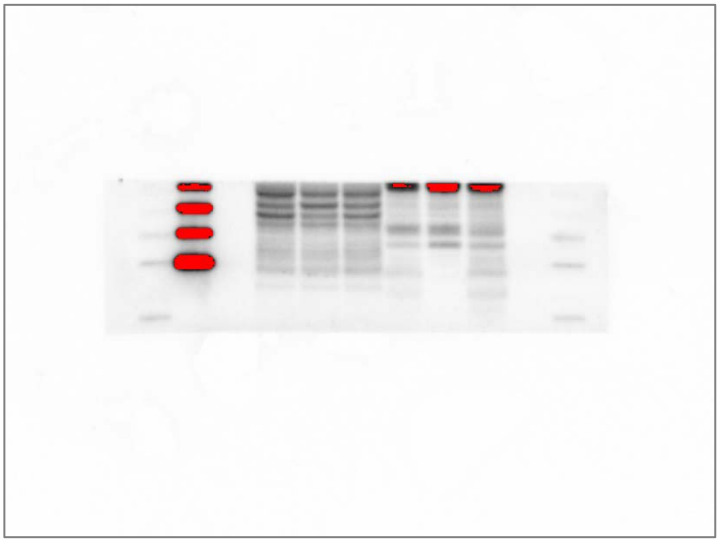

b

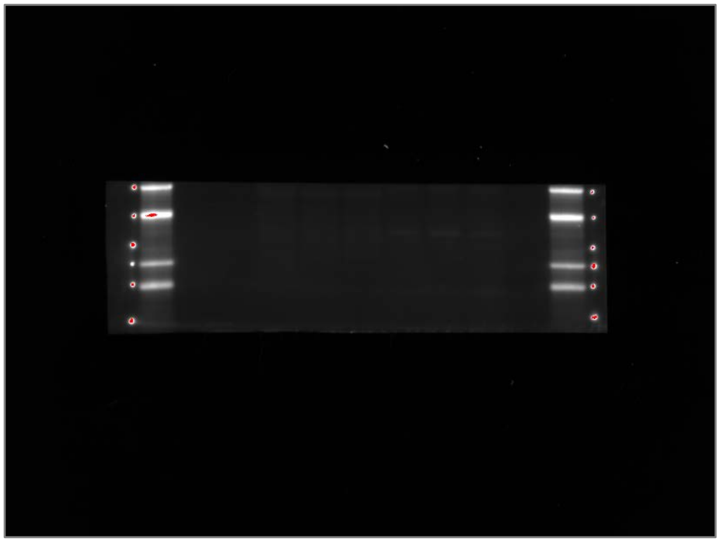

**Supplementary Fig. 12: Unedited western blot strip of CD81 (reduced conditions).**

Columns from left-to-right: Kaleidoscope Protein Standards, MagicMark Protein Standards, Empty Lane, 50 μg Clone 9 WCL, 50 μg RFL-6 WCL, 50 μg RMC WCL, 50 μg Clone 9 EV, 50 μg RFL-6 EV, 50 μg RMC EV, Empty Lane, Kaleidoscope Protein Standards.

For (a) chemiluminescent detection of HRP, image acquisition was performed using an exposure time of 4 seconds. Fluorescent detection (b) of the Kaleidoscope Protein Standards was performed using settings for Alexa Fluor 680 and optimized for intense bands. Red color represents signal saturation. Dots next to protein standards in the fluorescent image are a byproduct of labeling the visible standards with a laboratory marker. Western blot was cut just above the Kaleidoscope 50 kDa protein standard (top band in the fluorescent channel). MagicMark XP protein standards from top-down are: 50 kDa, 40 kDa, 30 kDa, 20 kDa.

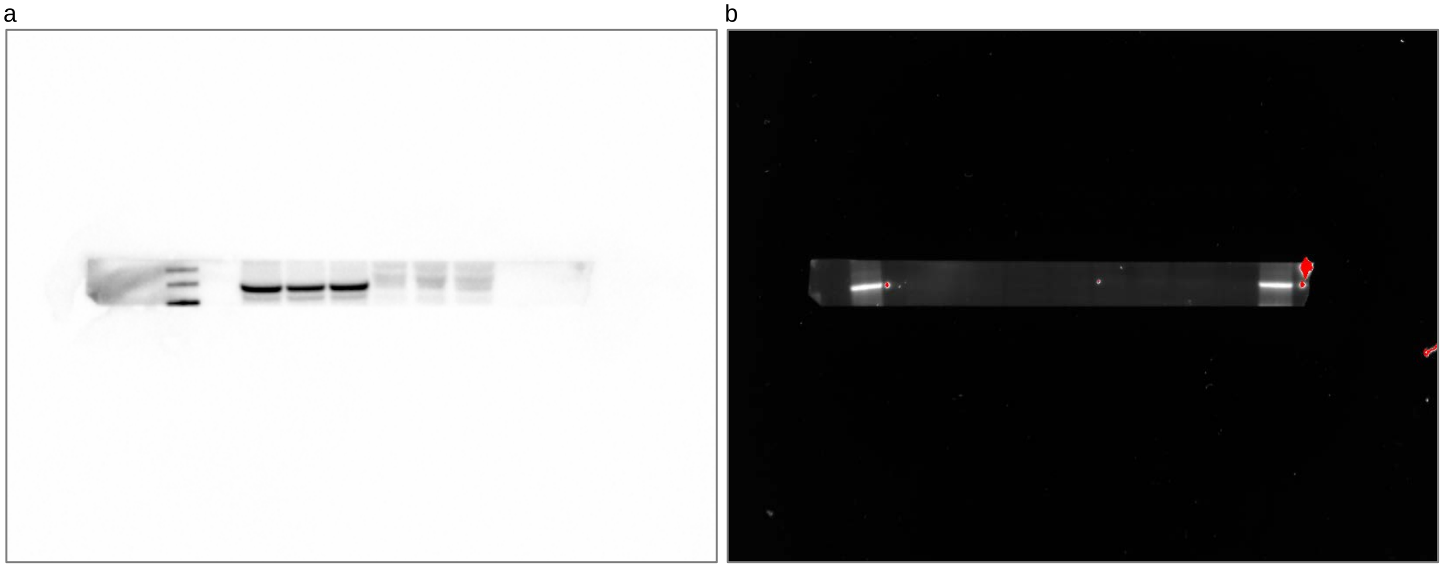

**Supplementary Fig. 13: Unedited western blot of tsg 101.**

Columns from left-to-right: Kaleidoscope Protein Standards, MagicMark Protein Standards, Empty Lane, 50 μg Clone 9 WCL, 50 μg RFL-6 WCL, 50 μg RMC WCL, 50 μg Clone 9 EV, 50 μg RFL-6 EV, 50 μg RMC EV, Empty Lane, Kaleidoscope Protein Standards.

For (a) chemiluminescent detection of HRP, image acquisition was performed using optimization for intense bands. Fluorescent detection (b) of the Kaleidoscope Protein Standards was performed using settings for Alexa Fluor 680 and optimized for intense bands. Red color represents signal saturation. Dots next to protein standards in the fluorescent image are a byproduct of labeling the visible standards with a laboratory marker. Western blot was cut above the Kaleidoscope 37 kDa protein standard and below the 75 kDa protein standard (not visible in the fluorescent channel). MagicMark XP protein standards from top-down are: 60 kDa, 50 kDa, 40 kDa.

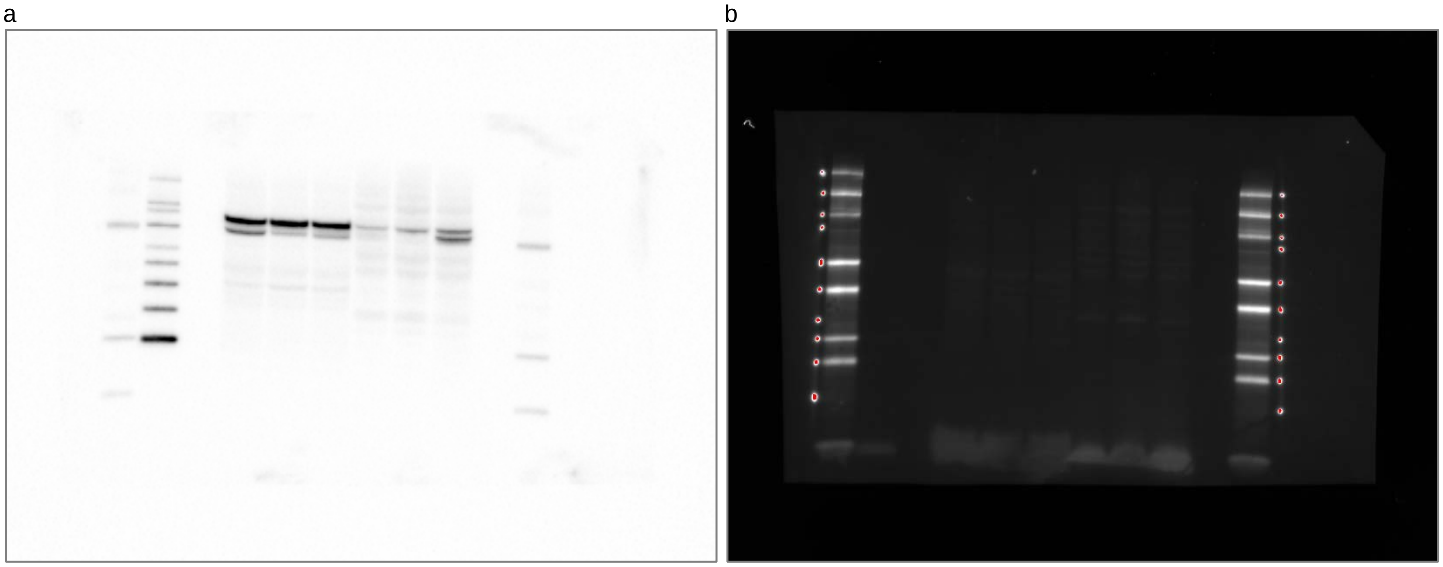

**Supplementary Fig. 14: Unedited western blot of Alix.**

Columns from left-to-right: Kaleidoscope Protein Standards, MagicMark Protein Standards, Empty Lane, 50 μg Clone 9 WCL, 50 μg RFL-6 WCL, 50 μg RMC WCL, 50 μg Clone 9 EV, 50 μg RFL-6 EV, 50 μg RMC EV, Empty Lane, Kaleidoscope Protein Standards.

For (a) chemiluminescent detection of HRP, image acquisition was performed using an exposure time of 0.18 seconds. Fluorescent detection (b) of the Kaleidoscope Protein Standards was performed using settings for Alexa Fluor 680 and optimized for intense bands. Red color represents signal saturation. Dots next to protein standards in the fluorescent image are a byproduct of labeling the visible standards with a laboratory marker.

a

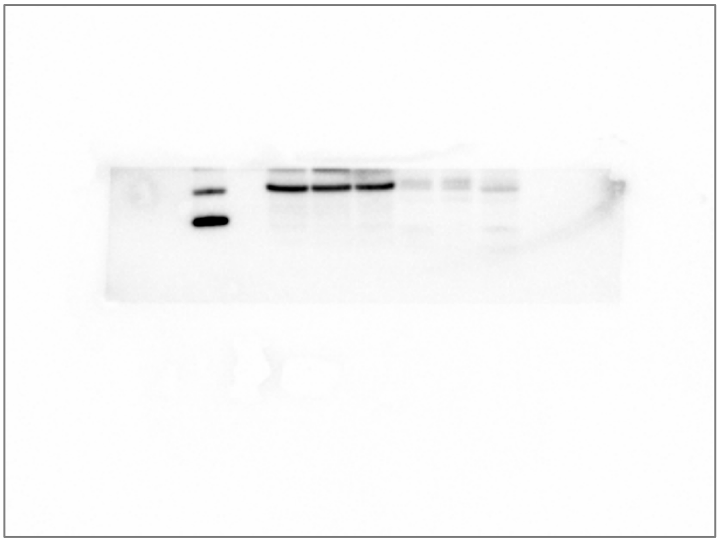

b

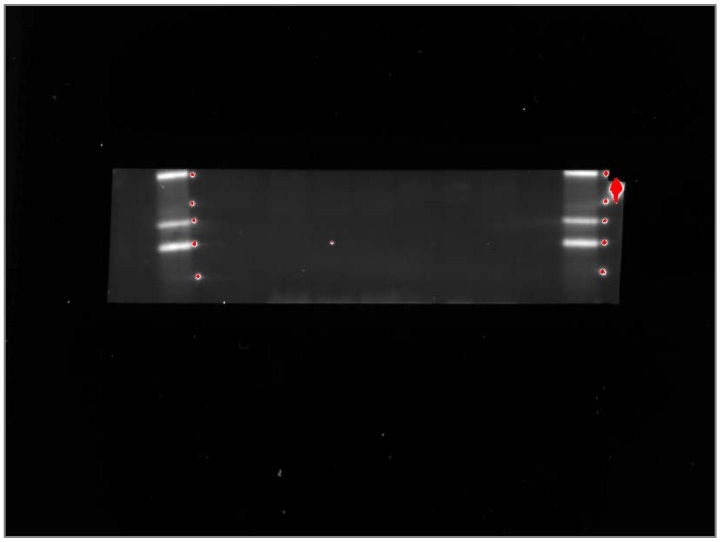

**Supplementary Fig. 15: Unedited western blot of ApoA-I.**

Columns from left-to-right: Kaleidoscope Protein Standards, MagicMark Protein Standards, Empty Lane, 50 μg Clone 9 WCL, 50 μg RFL-6 WCL, 50 μg RMC WCL, 50 μg Clone 9 EV, 50 μg RFL-6 EV, 50 μg RMC EV, Empty Lane, Kaleidoscope Protein Standards.

For (a) chemiluminescent detection of HRP, image acquisition was performed using optimization for intense bands. Fluorescent detection (b) of the Kaleidoscope Protein Standards was performed using settings for Alexa Fluor 680 and optimized for intense bands. Red color represents signal saturation. Dots next to protein standards in the fluorescent image are a byproduct of labeling the visible standards with a laboratory marker. Western blot was cut above the Kaleidoscope 37 kDa protein standard. MagicMark XP protein standards from top-down are: 40 kDa (half-visible), 30 kDa, 20 kDa.

a b

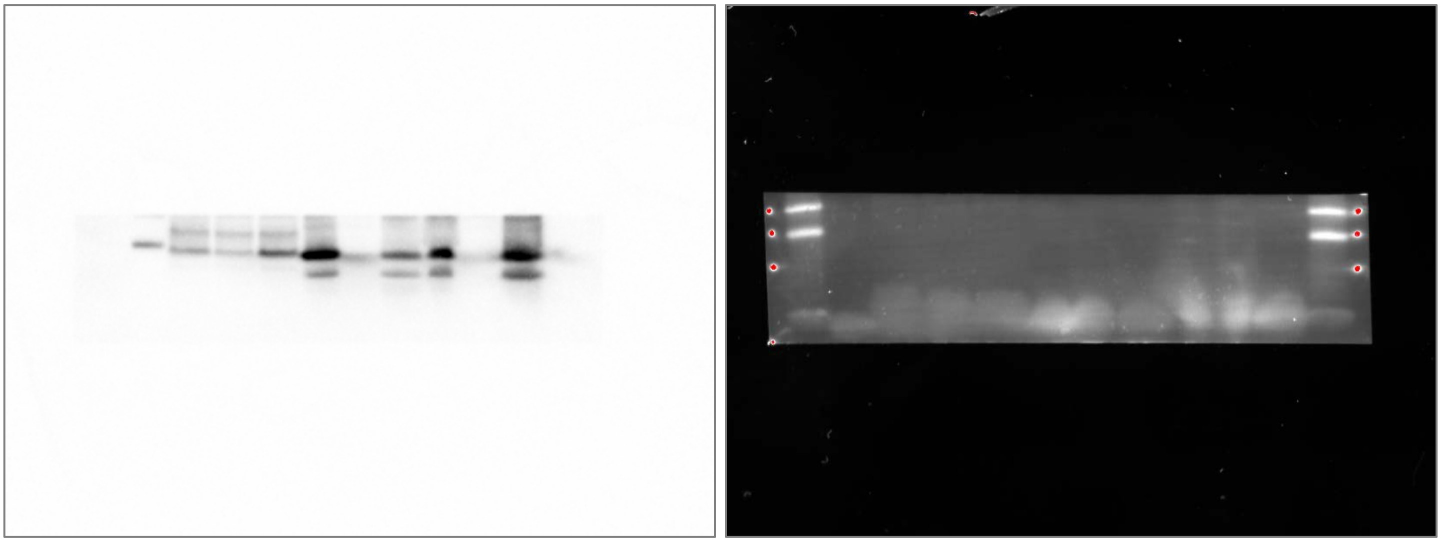

**Supplementary Fig. 16: Unedited western blot of Histone H3.1.**

Columns from left-to-right: Kaleidoscope Protein Standards, MagicMark Protein Standards, 50 μg Clone 9 WCL, 50 μg RFL-6 WCL, 50 μg RMC WCL, 50 μg Clone 9 EV, 50 μg RFL-6 EV, 50 μg RMC EV, 100 μg Clone 9 EV, 100 μg RFL-6 EV, 100 μg RMC EV, Kaleidoscope Protein Standards.

For (a) chemiluminescent detection of HRP, image acquisition was performed using optimization for intense bands. Fluorescent detection (b) of the Kaleidoscope Protein Standards was performed using settings for Alexa Fluor 680 and optimized for intense bands. Red color represents signal saturation. Dots next to protein standards in the fluorescent image are a byproduct of labeling the visible standards with a laboratory marker. Western blot was cut just below the Kaleidoscope 25 kDa protein standard. MagicMark XP protein standard is 20 kDa. Kaleidoscope protein standards from top-down: 20 kDa, 15 kDa, and 10 kDa (not visible but marked).

a

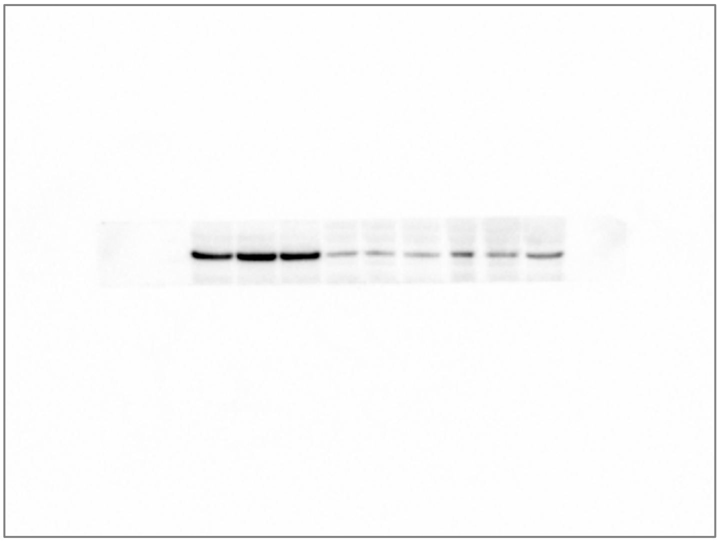

b

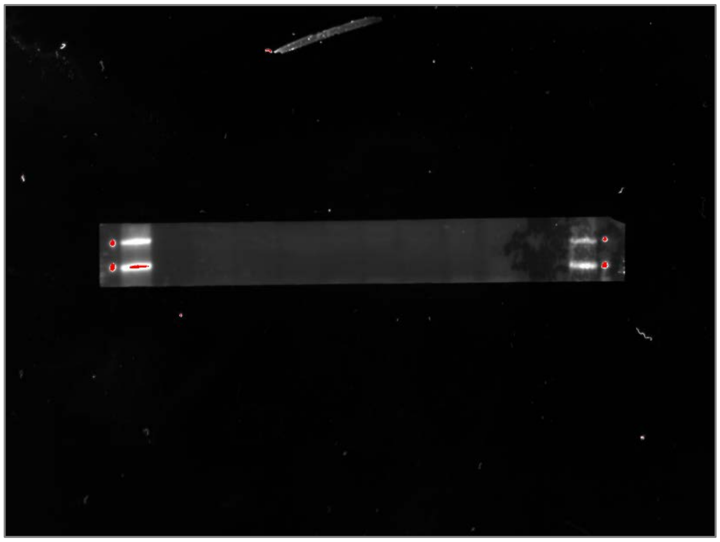

**Supplementary Fig. 17: Unedited western blot of Cyt c.**

Columns from left-to-right: Kaleidoscope Protein Standards, MagicMark Protein Standards, 50 μg Clone 9 WCL, 50 μg RFL-6 WCL, 50 μg RMC WCL, 50 μg Clone 9 EV, 50 μg RFL-6 EV, 50 μg RMC EV, 100 μg Clone 9 EV, 100 μg RFL-6 EV, 100 μg RMC EV, Kaleidoscope Protein Standards.

For (a) chemiluminescent detection of HRP, image acquisition was performed using optimization for intense bands. Fluorescent detection (b) of the Kaleidoscope Protein Standards was performed using settings for Alexa Fluor 680 and optimized for intense bands. Red color represents signal saturation. Dots next to protein standards in the fluorescent image are a byproduct of labeling the visible standards with a laboratory marker. Western blot was cut above the Kaleidoscope 50 kDa protein standard and below the 37 kDa protein standard. MagicMark XP protein standards were not visible at this exposure. Kaleidoscope protein standards from top-down are: 50 kDa, 37 kDa.

a b

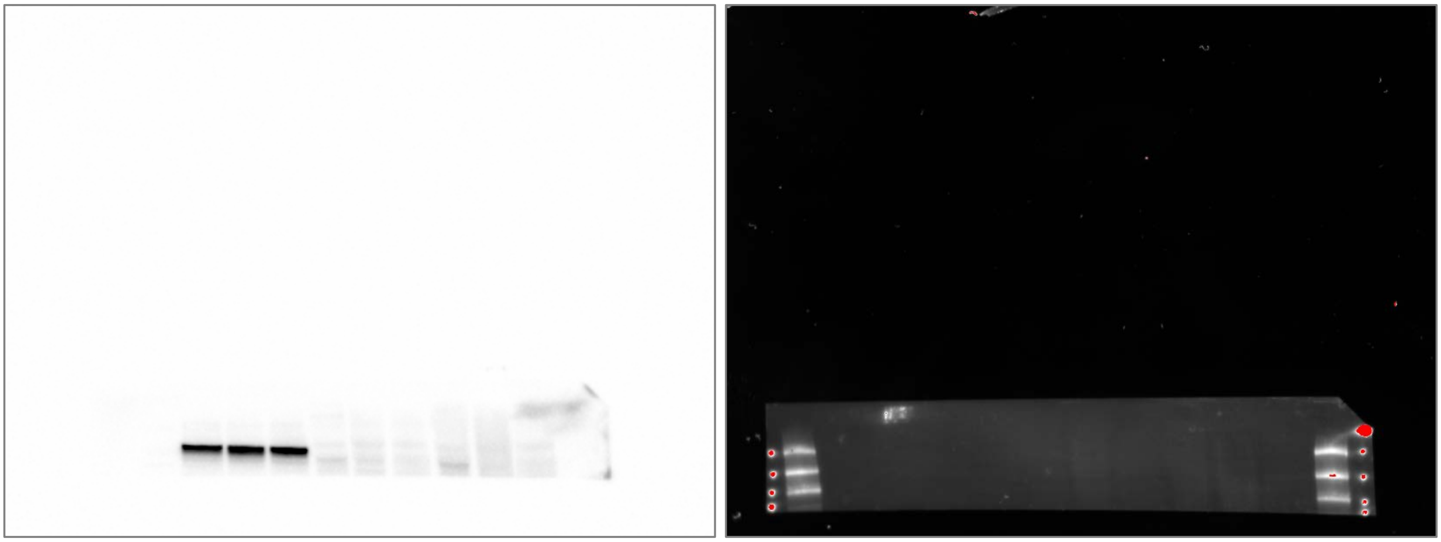

**Supplementary Fig. 18: Unedited western blot of GM130.**

Columns from left-to-right: Kaleidoscope Protein Standards, MagicMark Protein Standards, 50 μg Clone 9 WCL, 50 μg RFL-6 WCL, 50 μg RMC WCL, 50 μg Clone 9 EV, 50 μg RFL-6 EV, 50 μg RMC EV, 100 μg Clone 9 EV, 100 μg RFL-6 EV, 100 μg RMC EV, Kaleidoscope Protein Standards.

For (a) chemiluminescent detection of HRP, image acquisition was performed using optimization for intense bands. Fluorescent detection (b) of the Kaleidoscope Protein Standards was performed using settings for Alexa Fluor 680 and optimized for intense bands. Red color represents signal saturation. Dots next to protein standards in the fluorescent image are a byproduct of labeling the visible standards with a laboratory marker. Western blot was cut at the Kaleidoscope 75 kDa protein standard. MagicMark XP protein standards were not visible at this exposure. Kaleidoscope protein standards from top-down are: 250 kDa, 150 kDa, 100 kDa, and 75 kDa (half-visible in column 1).

a b

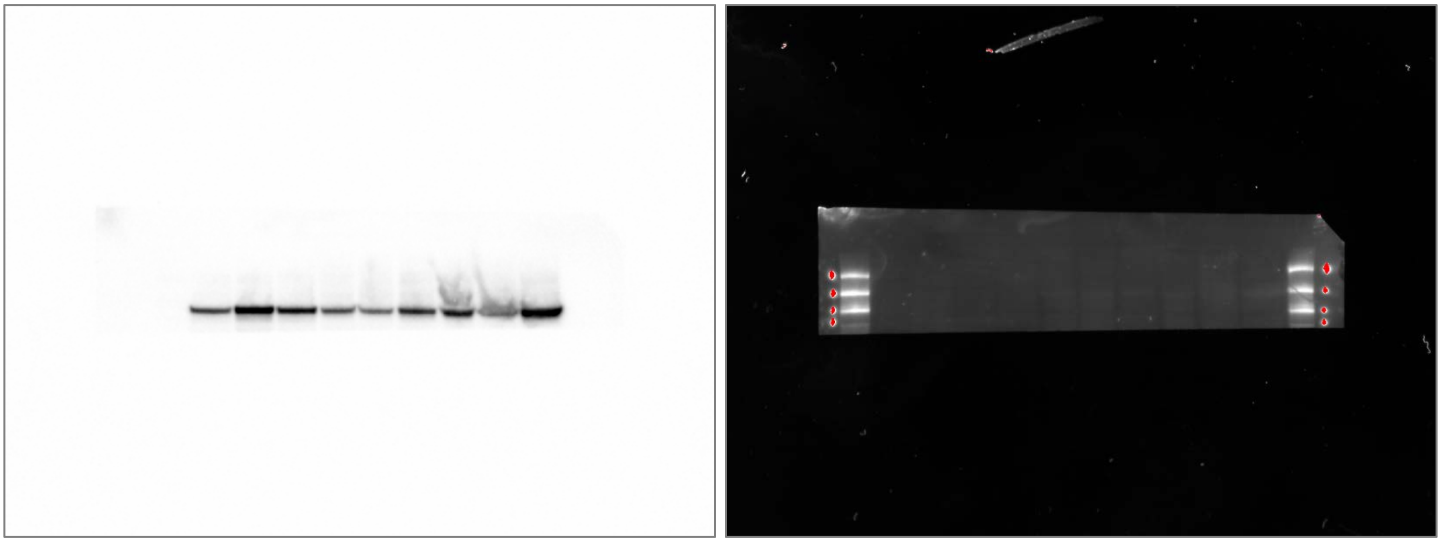

**Supplementary Fig. 19: Unedited western blot of α-actinin.**

Columns from left-to-right: Kaleidoscope Protein Standards, MagicMark Protein Standards, 50 μg Clone 9 WCL, 50 μg RFL-6 WCL, 50 μg RMC WCL, 50 μg Clone 9 EV, 50 μg RFL-6 EV, 50 μg RMC EV, 100 μg Clone 9 EV, 100 μg RFL-6 EV, 100 μg RMC EV, Kaleidoscope Protein Standards.

For (a) chemiluminescent detection of HRP, image acquisition was performed using optimization for intense bands. Fluorescent detection (b) of the Kaleidoscope Protein Standards was performed using settings for Alexa Fluor 680 and optimized for intense bands. Red color represents signal saturation. Dots next to protein standards in the fluorescent image are a byproduct of labeling the visible standards with a laboratory marker. Western blot was below the Kaleidoscope 75 kDa protein standard. MagicMark XP protein standards were not visible at this exposure. Kaleidoscope protein standards from top-down are: 250 kDa, 150 kDa, 100 kDa, and 75 kDa.

a b

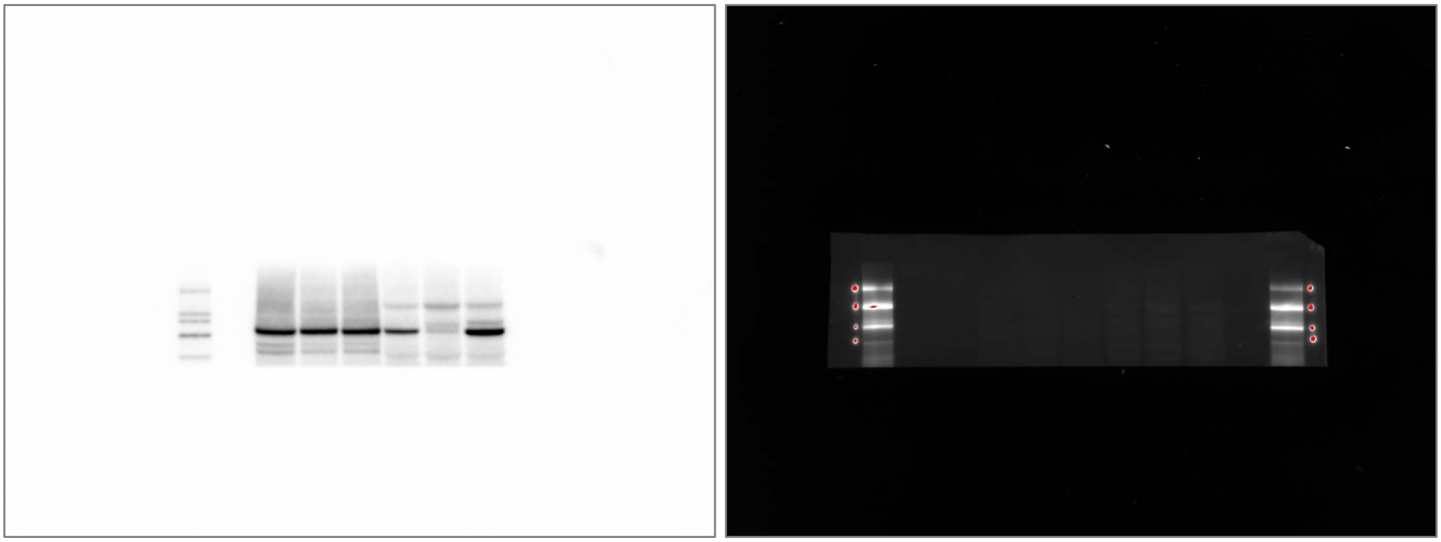

**Supplementary Fig. 20: Unedited western blot of Ago1-4.**

Columns from left-to-right: Kaleidoscope Protein Standards, MagicMark Protein Standards, Empty Lane, 50 μg Clone 9 WCL, 50 μg RFL-6 WCL, 50 μg RMC WCL, 50 μg Clone 9 EV, 50 μg RFL-6 EV, 50 μg RMC EV, Empty Lane, Kaleidoscope Protein Standards.

For (a) chemiluminescent detection of HRP, image acquisition was performed using optimization for intense bands. Fluorescent detection (b) of the Kaleidoscope Protein Standards was performed using settings for Alexa Fluor 680 and optimized for intense bands. Red color represents signal saturation. Dots next to protein standards in the fluorescent image are a byproduct of labeling the visible standards with a laboratory marker. Western blot was above the Kaleidoscope 50 kDa protein standard. MagicMark XP protein standards from top-down are: 220 kDa, 120 kDa, 100 kDa, 80 kDa, 60 kDa.

a

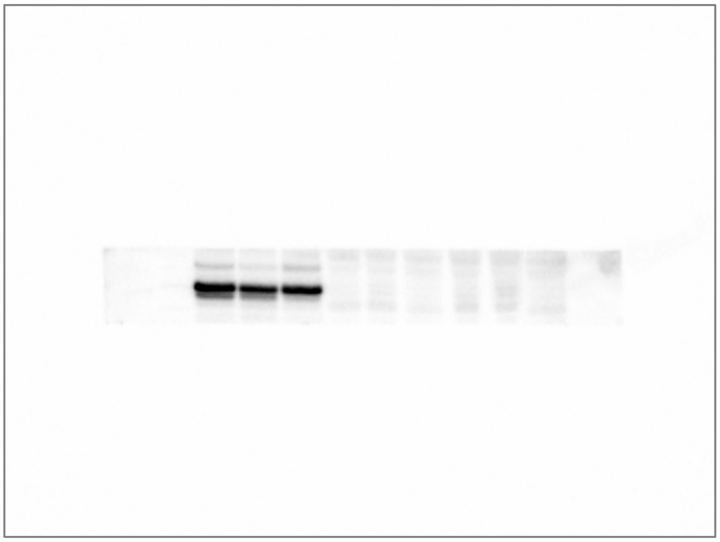

b

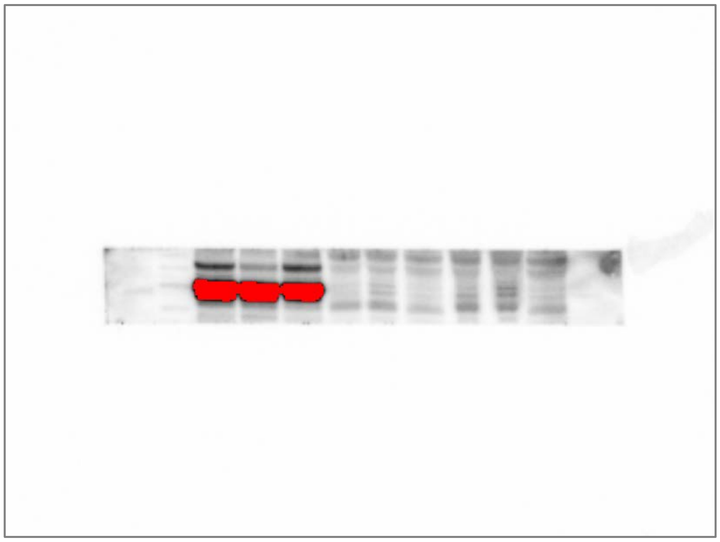

c

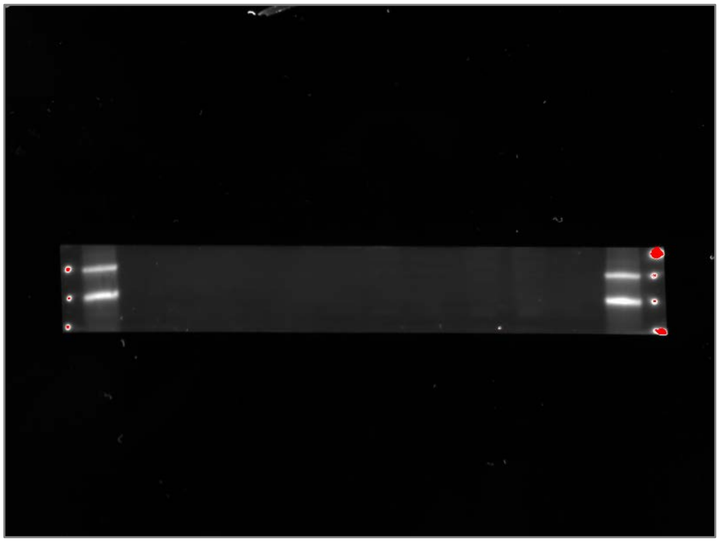

**Supplementary Fig. 21: Unedited western blot of hnRNP A2/B1.**

Columns from left-to-right: Kaleidoscope Protein Standards, MagicMark Protein Standards, 50 μg Clone 9 WCL, 50 μg RFL-6 WCL, 50 μg RMC WCL, 50 μg Clone 9 EV, 50 μg RFL-6 EV, 50 μg RMC EV, 100 μg Clone 9 EV, 100 μg RFL-6 EV, 100 μg RMC EV, Kaleidoscope Protein Standards.

For chemiluminescent detection of HRP, image acquisition was optimized for (a) intense bands and (b) faint bands. Fluorescent detection (c) of the Kaleidoscope Protein Standards was performed using settings for Alexa Fluor 680 and optimized for intense bands. Red color represents signal saturation. Dots next to protein standards in the fluorescent image are a byproduct of labeling the visible standards with a laboratory marker. Western blot was cut just below the Kaleidoscope 75 kDa protein standard and just below the 25 kDa protein standard. MagicMark XP protein standards in (b) from top-down are: 60 kDa, 50 kDa, 40 kDa, 30 kDa. Kaleidoscope protein standards from top-down are: 50 kDa, 37 kDa, and 25 kDa (not visible but marked).
