## Supplementary Note for "Quantification of extracellular vesicles with unaltered surface membranes using an internalized oligonucleotide tracer and droplet digital PCR applied to modeling of *in vivo* kinetics"

**Table of Contents**


1. 3 compartment with covariates model code
2. 3 compartment with covariates model code executed

3 compartment with covariates model code:

```
test(){
	cfMicro(A1, Cl / V, Cl2 / V, Cl2 / V2, Cl3 / V, Cl3 / V3)
	dosepoint(A1)
	C = A1 / V
	error(CEps = 0.448343)
	observe(CObs = C * (1 + CEps))
	stparm(V = tvV * exp(nV))
	stparm(V2 = tvV2 * exp(dV2dCellLinenum1*(CellLinenum==1)) *
exp(dV2dCellLinenum2*(CellLinenum==2)) * exp(nV2))
	stparm(V3 = tvV3 * exp(dV3dCellLinenum1*(CellLinenum==1)) *
exp(dV3dCellLinenum2*(CellLinenum==2)) * exp(nV3))
	stparm(Cl = tvCl * exp(dCldCellLinenum1*(CellLinenum==1)) *
exp(dCldCellLinenum2*(CellLinenum==2)) * exp(nCl))
	stparm(Cl2 = tvCl2 * exp(nCl2))
	stparm(Cl3 = tvCl3 * exp(dCl3dCellLinenum1*(CellLinenum==1)) *
exp(dCl3dCellLinenum2*(CellLinenum==2)) * exp(nCl3))
	fcovariate(CellLinenum())
	fixef(tvV = c(, 27669.7, ))
	fixef(tvV2 = c(, 3060570, ))
	fixef(tvV3 = c(, 16001.1, ))
	fixef(tvCl = c(, 93445, ))
	fixef(tvCl2 = c(, 111078, ))
	fixef(tvCl3 = c(, 21211.7, ))
	fixef(dV2dCellLinenum1(enable=c(0)) = c(, -0.0247024, ))
	fixef(dV2dCellLinenum2(enable=c(0)) = c(, -1.40195, ))
	fixef(dV3dCellLinenum1(enable=c(1)) = c(, 1.72244, ))
	fixef(dV3dCellLinenum2(enable=c(1)) = c(, 0.077768, ))
	fixef(dCldCellLinenum1(enable=c(2)) = c(, -1.57004, ))
	fixef(dCldCellLinenum2(enable=c(2)) = c(, -0.34285, ))
	fixef(dCl3dCellLinenum1(enable=c(3)) = c(, 1.76805, ))
	fixef(dCl3dCellLinenum2(enable=c(3)) = c(, 1.09156, ))
	ranef(diag(nV, nV2, nCl, nCl2, nV3, nCl3) = c(3.3924198E-05, 0.87921569, 0.76057715,
0.24396838, 0.0047016738, 5.3406114E-05))
}
```

```
3 compartment with covariates model code executed:

# Population:
 ELS FOCE/general Laplace engine log file

 MAX major iterations =          10
 MAX Niter (line searches per major iteration) =        1000

 ---------NEW MAJOR ITERATION--------
 Major iteration =           1

 FOCE Hessian approximation used

 Model evaluation ODE level used:
 0=none

 Other flags and run conditions/tolerances:

        28  NSUB
        21  NUMFREEPARAM
        14  NUMFREEFIXEF
         1  NUMFREEEPS
         6  NUMFREEOMEGAPARAM
         6  NUMRANEF
         1  Allow Gaussian Fit
         5  IDENGINE
         0  IFLAGRESTART
         1  IFLAGSTDERR
         0  IFLAGNP
         1  IFLAGFOCEHESS
         1  NORDERAGQ
         1  NUMPROCS
        13  NDIGITblup
         7  NDIGITlagl
 0.200E-02  tolmodlinz
 0.100E-01  tolstderr
       100  NREP_PCWRES
         1  NPRESAMPLE
         0  NGETMAPNP
           0  iflaganagrad
           0  iodelevelused
           1  iflagstderr
           1  iflagwhichstderr


 Initial parameter values:

  0.27669700E+05        1   1 THETA
  0.30605700E+07        2   1 THETA
  0.16001100E+05        3   1 THETA
  0.93445000E+05        4   1 THETA
  0.11107800E+06        5   1 THETA
  0.21211700E+05        6   1 THETA
 -0.24702400E-01        7   1 THETA
 -0.14019500E+01        8   1 THETA
  0.17224400E+01        9   1 THETA
  0.77768000E-01       10   1 THETA
 -0.15700400E+01       11   1 THETA
 -0.34285000E+00       12   1 THETA
  0.17680500E+01       13   1 THETA
  0.10915600E+01       14   1 THETA
  0.44834300E+00        1   1 EPS_STD_DEV
  0.20101145E+00        1   1 EPS_VARIANCE
  0.33924198E-04        1   1 OMEGA
  0.00000000E+00        2   1 OMEGA
  0.87921569E+00        2   2 OMEGA
  0.00000000E+00        3   1 OMEGA
```

```
 0.00000000E+00        3   2 OMEGA
 0.76057715E+00        3   3 OMEGA
 0.00000000E+00        4   1 OMEGA
 0.00000000E+00        4   2 OMEGA
 0.00000000E+00        4   3 OMEGA
 0.24396838E+00        4   4 OMEGA
 0.00000000E+00        5   1 OMEGA
 0.00000000E+00        5   2 OMEGA
 0.00000000E+00        5   3 OMEGA
 0.00000000E+00        5   4 OMEGA
 0.47016738E-02        5   5 OMEGA
 0.00000000E+00        6   1 OMEGA
 0.00000000E+00        6   2 OMEGA
 0.00000000E+00        6   3 OMEGA
 0.00000000E+00        6   4 OMEGA
 0.00000000E+00        6   5 OMEGA
 0.53406114E-04        6   6 OMEGA
-0.17046089E+04        1   1 -LOGLIKE
-0.39219854E+04        1   1 ELSOBJ

Total # Subjects     =          28
Total # Observations =         279


-loglike at initial solution
after initial stabilization =  -1704.6088625882091

Main OPTIF9 optimization for engine 5 terminated
ITRMCD exit code from OPTIF9 =           1
1,2,3=probable success,4=maxiterations reached


Starting std error computation

-2LL Hessian inverse std error successful
   i   internal par(i)   std. error      rel. std. error

   1    0.276831E+05    0.223182E+04        0.080620
   2    0.305655E+07    0.815136E+06        0.266685
   3    0.160072E+05    0.408402E+04        0.255136
   4    0.934983E+05    0.238597E+05        0.255188
   5    0.111156E+06    0.133762E+05        0.120337
   6    0.212235E+05    0.421109E+04        0.198417
   7   -0.247059E-01    0.114009E-01        0.461466
   8   -0.140210E+01    0.381477E+00        0.272075
   9    0.172296E+01    0.270328E+00        0.156898
  10    0.777273E-01    0.341233E-01        0.439013
  11   -0.156984E+01    0.442076E+00        0.281605
  12   -0.342931E+00    0.151701E+00        0.442364
  13    0.176776E+01    0.247251E+00        0.139867
  14    0.109080E+01    0.394064E+00        0.361263
  15    0.448256E+00    0.271243E-01        0.060511
  16    0.582394E-02    0.377574E-02        0.648314
  17    0.937732E+00    0.160040E+00        0.170668
  18    0.872024E+00    0.144851E+00        0.166109
  19    0.493714E+00    0.886871E-01        0.179633
  20    0.685832E-01    0.435955E-01        0.635659
  21    0.730711E-02    0.512304E-02        0.701103


Standard errors of estimated parameters

internal_coords    Param_val       Hessinv         Sandwich        Score

         1        0.276831E+05    0.223182E+04    0.192395E+04    0.673319E+04
         2        0.305655E+07    0.815136E+06    0.866155E+06    0.239797E+07
         3        0.160072E+05    0.408402E+04    0.624424E+04    0.475476E+04
```

```
         4        0.934983E+05    0.238597E+05    0.192119E+05    0.805001E+05
         5        0.111156E+06    0.133762E+05    0.159945E+05    0.186205E+05
         6        0.212235E+05    0.421109E+04    0.311962E+04    0.101803E+05
         7       -0.247059E-01    0.114009E-01    0.118292E-02    0.596376E+00
         8       -0.140210E+01    0.381477E+00    0.354022E+00    0.123236E+01
         9        0.172296E+01    0.270328E+00    0.383587E+00    0.326966E+00
        10        0.777273E-01    0.341233E-01    0.583719E-02    0.945907E+00
        11       -0.156984E+01    0.442076E+00    0.459944E+00    0.115293E+01
        12       -0.342931E+00    0.151701E+00    0.635240E-01    0.122391E+01
        13        0.176776E+01    0.247251E+00    0.196924E+00    0.523298E+00
        14        0.109080E+01    0.394064E+00    0.419760E+00    0.989361E+00
        15        0.448256E+00    0.271243E-01    0.336418E-01    0.381471E-01
        16        0.582394E-02    0.377574E-02    0.838928E-03    0.243198E+00
        17        0.937732E+00    0.160040E+00    0.185836E+00    0.382713E+00
        18        0.872024E+00    0.144851E+00    0.152279E+00    0.381116E+00
        19        0.493714E+00    0.886871E-01    0.769431E-01    0.215328E+00
        20        0.685832E-01    0.435955E-01    0.132987E-01    0.833214E+00
        21        0.730711E-02    0.512304E-02    0.214727E-02    0.107418E+00

external_coords    Param_val       Hessinv         Sandwich        Score

         1        0.276831E+05    0.223182E+04    0.192395E+04    0.673319E+04
         2        0.305655E+07    0.815136E+06    0.866155E+06    0.239797E+07
         3        0.160072E+05    0.408402E+04    0.624424E+04    0.475476E+04
         4        0.934983E+05    0.238597E+05    0.192119E+05    0.805001E+05
         5        0.111156E+06    0.133762E+05    0.159945E+05    0.186205E+05
         6        0.212235E+05    0.421109E+04    0.311962E+04    0.101803E+05
         7       -0.247059E-01    0.114009E-01    0.118292E-02    0.596376E+00
         8       -0.140210E+01    0.381477E+00    0.354022E+00    0.123236E+01
         9        0.172296E+01    0.270328E+00    0.383587E+00    0.326966E+00
        10        0.777273E-01    0.341233E-01    0.583719E-02    0.945907E+00
        11       -0.156984E+01    0.442076E+00    0.459944E+00    0.115293E+01
        12       -0.342931E+00    0.151701E+00    0.635240E-01    0.122391E+01
        13        0.176776E+01    0.247251E+00    0.196924E+00    0.523298E+00
        14        0.109080E+01    0.394064E+00    0.419760E+00    0.989361E+00
        15        0.448256E+00    0.271243E-01    0.336418E-01    0.381471E-01
        16        0.339182E-04    0.439794E-04    0.977173E-05    0.283274E-02
        17        0.879341E+00    0.300150E+00    0.348528E+00    0.717764E+00
        18        0.760425E+00    0.252626E+00    0.265581E+00    0.664684E+00
        19        0.243753E+00    0.875720E-01    0.759757E-01    0.212621E+00
        20        0.470366E-02    0.597984E-02    0.182414E-02    0.114289E+00
        21        0.533939E-04    0.748693E-04    0.313806E-04    0.156984E-02

Engine Convergence Summary:
Convergence achieved

OPTIF9 ITRMCD value on last iteration =           1
NLME7 return code=           1

Optimal parameters:

 0.27683136E+05        1   1 THETA        BOUNDS:  -0.100+101  0.100+101
 0.30565508E+07        2   1 THETA        BOUNDS:  -0.100+101  0.100+101
 0.16007238E+05        3   1 THETA        BOUNDS:  -0.100+101  0.100+101
 0.93498288E+05        4   1 THETA        BOUNDS:  -0.100+101  0.100+101
 0.11115596E+06        5   1 THETA        BOUNDS:  -0.100+101  0.100+101
 0.21223484E+05        6   1 THETA        BOUNDS:  -0.100+101  0.100+101
-0.24705900E-01        7   1 THETA        BOUNDS:  -0.100+101  0.100+101
-0.14021015E+01        8   1 THETA        BOUNDS:  -0.100+101  0.100+101
 0.17229560E+01        9   1 THETA        BOUNDS:  -0.100+101  0.100+101
 0.77727348E-01       10   1 THETA        BOUNDS:  -0.100+101  0.100+101
-0.15698417E+01       11   1 THETA        BOUNDS:  -0.100+101  0.100+101
-0.34293142E+00       12   1 THETA        BOUNDS:  -0.100+101  0.100+101
 0.17677641E+01       13   1 THETA        BOUNDS:  -0.100+101  0.100+101
 0.10907973E+01       14   1 THETA        BOUNDS:  -0.100+101  0.100+101
 0.44825609E+00        1   1 EPS_STD_DEV  BOUNDS:   0.100E-02  0.100+101
 0.20093352E+00        1   1 EPS_VARIANCE
```

```
 0.33918246E-04        1   1 OMEGA
 0.00000000E+00        2   1 OMEGA
 0.87934107E+00        2   2 OMEGA
 0.00000000E+00        3   1 OMEGA
 0.00000000E+00        3   2 OMEGA
 0.76042537E+00        3   3 OMEGA
 0.00000000E+00        4   1 OMEGA
 0.00000000E+00        4   2 OMEGA
 0.00000000E+00        4   3 OMEGA
 0.24375324E+00        4   4 OMEGA
 0.00000000E+00        5   1 OMEGA
 0.00000000E+00        5   2 OMEGA
 0.00000000E+00        5   3 OMEGA
 0.00000000E+00        5   4 OMEGA
 0.47036575E-02        5   5 OMEGA
 0.00000000E+00        6   1 OMEGA
 0.00000000E+00        6   2 OMEGA
 0.00000000E+00        6   3 OMEGA
 0.00000000E+00        6   4 OMEGA
 0.00000000E+00        6   5 OMEGA
 0.53393923E-04        6   6 OMEGA
-0.17046094E+04        1   1 -LOGLIKE
-0.39219865E+04        1   1 ELSOBJ
 engine runtime (secs) =           1.109
 stderr runtime (secs) =           6.844


SHRINKAGES

  eps-shrinkage     =          0.11931

  eta-shrinkage( 1) =          0.98788
  eta-shrinkage( 2) =          0.12678
  eta-shrinkage( 3) =          0.14688
  eta-shrinkage( 4) =          0.13362
  eta-shrinkage( 5) =          0.87521
  eta-shrinkage( 6) =          0.98909

RUNTIMES

  engine runtime (secs) =      1.109
  stderr runtime (secs) =      6.844

EXITCODE            1
```
